# ROCKETS - a novel one-for-all toolbox for light sheet microscopy in drug discovery

**DOI:** 10.1101/2022.09.26.509149

**Authors:** Joerg PJ Mueller, Michael Dobosz, Nils O’Brien, Anna Maria Giusti, Martin Lechmann, Franz Osl, Ann-Katrin Wolf, Markus Sauer, Frank Herting, Pablo Umana, Sara Colombetti, Thomas Pöschinger, Andreas Beilhack

**Affiliations:** Interdisciplinary Center for Clinical Research Laboratory (IZKF) Würzburg, Department of Internal Medicine II, Center for Experimental Molecular Medicine, Würzburg University Hospital, Würzburg, Germany; Roche Pharmaceutical Research and Early Development, Roche Diagnostics GmbH, Nonnenwald 2, Penzberg, Germany; Roche Pharmaceutical Research and Early Development, Roche Glycart AG, Wagistraße 10, Schlieren, Switzerland; Department of Biotechnology and Biophysics, Biocenter, University of Würzburg, Würzburg, Germany; Regeneron Pharmaceuticals, New York

**Keywords:** Imaging, immunotherapy, preclinical drug development, biodistribution, cancer, light sheet fluorescence microscopy, EpCAM

## Abstract

Advancing novel immunotherapy strategies requires refined tools in preclinical research to thoroughly assess drug targets, biodistribution, safety, and efficacy. Light sheet fluorescence microscopy (LSFM) offers unprecedented fast volumetric ex vivo imaging of large tissue samples in high resolution. Yet, to date laborious and unstandardized tissue processing procedures have limited throughput and broader applications in immunological research. Therefore, we have developed a simple and harmonized protocol for processing, clearing and imaging of all mouse organs and even entire mouse bodies. Applying this Rapid Optical Clearing Kit for Enhanced Tissue Scanning (ROCKETS) in combination with LSFM allowed us to comprehensively study the in vivo biodistribution of an antibody targeting Epithelial Cell Adhesion Molecule (EpCAM) in 3D. Quantitative high-resolution scans of whole organs did not only reveal known EpCAM expression patterns but, importantly, uncovered several new EpCAM-binding sites. We identified choroid plexi in the brain and duodenal papillae as unexpected locations of high EpCAM-expression. These tissue locations may be considered as particularly sensitive sites due their importance for liquor production or as critical junctions draining bile and digestive pancreatic enzymes into the small bowel, respectively. These newly gained insights appear highly relevant for clinical translation of EpCAM-addressing immunotherapies. Thus, ROCKETS in combination with LSFM may help to set new standards for preclinical evaluation of immunotherapeutic strategies. Conclusively, we propose ROCKETS as an ideal platform for a broader application of LSFM in immunological research optimally suited for quantitative co-localization studies of immunotherapeutic drugs and defined cell populations in the microanatomical context of organs or even whole mice.

## 1 Introduction

Paradigm shifting mechanistic insights, conceptual advances, and compelling clinical outcomes have placed immunotherapy at center stage in the treatment of cancer patients. Direct targeting of cancer cells with therapeutic monoclonal antibodies (1–3), T cell-engaging antibody formats (4), antibody-drug conjugates (5, 6) and radioimmunotherapy (7), genetically modified chimeric antigen-receptor (CAR) (8, 9) or T-cell receptor (TCR) T cells (10–12), and vaccination strategies (13–15) build an increasing armamentarium to treat cancer patients. Therapeutic approaches to indirectly boost the body’s natural defense against cancer have successfully improved clinical care by either targeting the cancer cells directly or the tumor microenvironment (3, 16) through blocking immune check points (12, 17, 18) and activating preexisting endogenous immune effector mechanisms (19–21). To date, clinical success has cemented immunotherapy as a powerful pillar of modern cancer therapy. Yet, directing and taming the powers of an effective immune response against cancer cells remains challenging (22–25). Clearly, insights in the spatial organization of cancer, stroma and immune cell topography, distribution and the molecular regulation of potential therapeutic target antigens and the local and systemic regulation of immune effector mechanisms are key aspects determining success or failure of novel therapeutic strategies. Preclinical development requires careful consideration of tumor and tissue antigens as well as complex heterogeneous tumor microenvironments. Identifying promising targets implies to subsequently outweighing therapeutic benefits with potential toxicities, which remains a major challenge during preclinical and clinical development.

EpCAM (CD326) was one of the first human tumor-associated antigens (TAA) discovered with monoclonal antibodies more than forty years ago in patients with colorectal carcinomas (26). Since then it has become clear that many solid cancers of epithelial origin, such as colon, breast, pancreas and prostate carcinomas, aberrantly overexpress EpCAM. EpCAM fulfills many functions in the regulation of cell adhesion, proliferation, migration, stemness, and epithelial-to-mesenchymal transition (EMT) of carcinoma cells (reviewed in (27)). Notably, many healthy epithelial tissues also express EpCAM. However, healthy simple and pseudostratified epithelia in humans express EpCAM in basolateral membranes with the exception of hepatocytes and keratinocytes in contrast to the ubiquitous non-polarized overexpression profile in epithelial cancer cells, (28, 29). These differential expression patterns have positioned EpCAM as an interesting antigen for targeted cancer therapy (27, 30) although EpCAM-targeted therapies must be closely assessed for on-target/off-tumor binding potentially resulting in adverse effects.

Therefore, advancing immunotherapies requires to further develop suitable tools and technologies to accelerate robust preclinical evaluation into successful clinical development.

Before entering clinical trials, any drug candidate must undergo extensive preclinical testing with the aim of predicting pharmacological properties and toxicological effects in humans (31). Herein, the *ex vivo* analysis of tissue specimens is often carried out based on histology. Modern histopathological analyses can rely on robust and highly standardized sample preparation techniques that have evolved over decades. However, it has been demonstrated that thin sections of embedded tissues are not always representative for the entire specimen (32, 33). Furthermore, creating hundreds of physical sections is extremely time consuming, laborious and uses up the specimen for further analysis, especially when rare events need to be detected within large tissue specimens (34–36).

Over the last two decades, light sheet fluorescence microscopy (LSFM) has emerged as a non-destructive technology offering rapid high-resolution imaging by creating optical sections of large intact tissue specimens (37). Consequently, LSFM has been applied across many fields of research (38) like developmental biology (39–43), neurobiology (44–47), cancer research (48–51) and immunology (52–56). High acquisition rates and recent progress towards batch-wise imaging of multiple specimens render LSFM principally suitable for large-scale preclinical studies with dozens or even hundreds of samples. However, as a prerequisite for mesoscopic LSFM imaging, specimens must be rendered optically transparent (clearing) (57–59). Clearing is generally achieved by removing light absorbing components from the tissue and reducing scattering through the homogenization of different refractive indices (RI) (58). Many protocols have emerged for the clearing of murine and human tissues, but most published procedures are limited to processing few specimens at a time and are often tailored for a specific tissue of interest (60, 61). For some tissues like the small and large intestine, no procedures exist that enable clearing of entire organs, rather than small segments (52, 62–65). Additionally, almost all protocols for murine tissues require animals to be perfused, a laborious and time-consuming procedure to flush out the blood from animals (66). Due to these limitations experimenters face great complexity if they want to clear more than one type of tissue or many specimens in parallel. Therefore, sample preparation still obstructs LSFM-based studies in preclinical drug development.

To this end we report three advances to overcome current challenges to routinely apply LSFM for advancing novel immunotherapy strategies. First, we combined and harmonized a clearing procedure of murine specimens optimally suited for standardized and high-throughput LSFM. Our *Rapid Optical Clearing Kit for Enhanced Tissue Scanning* (ROCKETS) approach, which does not require transcardial perfusion, combines in the first step hydrophilic expansion (hyperhydration), delipidation and decolorization and in the second step dehydration and organic solvent-based RI-matching. Second, we developed a technique for LSFM analysis of the entire gastrointestinal tract (GIT), which we termed 3D-Swiss Rolls technique. Third, we demonstrate that ROCKETS is also suited for LSFM of whole mouse bodies.

Finally, we investigated the biodistribution of an EpCAM-specific antibody employing ROCKETS and semiquantitative LSFM imaging, which resulted in unanticipated outcomes.

On top of confirming well recognized sites of EpCAM expression we report to our knowledge for the first time accentuated EpCAM-expression at all types of gustatory papillae of the tongue and especially in the choroid plexi in brain ventricles as well as the duodenal papillae. We deem the latter two unanticipated observations as highly relevant to be considered for cancer immunotherapeutic approaches. In summary we propose ROCKETS combined with LSFM to complement current immunohistochemical analyses for large scale assessment of in vivo drug development to advance immunotherapy.

## 2 Methods

### Animal models, handling and care

Female ten weeks old C57BL/6 inbred mice were obtained from Charles River Laboratories Germany GmbH, Sulzfeld, Germany. Studies were approved by the Government of Upper Bavaria (Regierung von Oberbayern, Munich, Germany) and in accordance with the European directive 2010/63/EU for animal research. For subcutaneous tumors, 3×10^5^ murine pancreatic ductal adenocarcinoma cells KPC-4662wt were applied as suspension in 100 µl matrigel (50% (v/v)), FisherScientific, Corning™ 354234) into the right flank of the animals. Animals were euthanized by cervical dislocation for whole organ analyses.

### Administration of conjugated antibodies

20 µg of anti-mouse EpCAM (CD326) antibody, conjugated with AlexaFluor750 (R&D Systems, FAB8998S, clone G8.8R), was administered to mice intravenously (i.v.) into the tail vein 24 h before euthanasia.

### Fixation of organs

Tissues of interest were excised immediately after euthanasia and rinsed briefly with deionized water (dH20) to remove hair or body fluids. Specimens were transferred to histological cassettes (Simport, Macrosette M512) for fixation. Tissues of the small and large intestines were processed according to the 3D-Swiss Roll procedure (below). All tissues were fixed using neutral buffered formaldehyde solution (NBF, 4% formaldehyde, VWR Chemicals 9713.9025) of at least ten times the volume of the dissected specimens for 14-18 h at 4°C with gentle agitation in the dark.

### Preclearing

For clearing of blood-rich and large mouse organs without transcardial perfusion, light absorbing and scattering tissue components were chemically removed by immersion in a preclearing reagent before dehydration and organic solvent-based RI-matching:

Fixed tissues were incubated at 30 °C in a minimum of 15 ml per whole organ with gentle agitation in the dark for two-four days, with one exchange after two days. The preclearing reagent comprised 20 % (v/v) Quadrol®, (N,N,N′,N′-Tetrakis(2-Hydroxypropyl)ethylenediamine, CAS102-60-3, Sigma Aldrich 122262-1L), 10 % (v/v) TWEEN-80® (Polyethylene glycol sorbitan monooleate, CAS 9005-65-6, Sigma Aldrich P1754-500ML), 10 % (v/v) TEA (2,2′,2′′-Nitrilotriethanol, CAS 102-71-6, Sigma Aldrich 90279), 10 % (v/v) DMSO (Dimethyl Sulfoxide, CAS 67-68-5, Sigma Aldrich D5879-500ML), 10 % (w/v) urea (CAS 57-13-6, Sigma Aldrich U5378-1KG) dissolved in dH2O. For mixing ∼100 ml/l of dH2O was applied before adding other components. Quadrol was heated up to ∼40°C to reduce viscosity and enable pouring.

After preclearing of organs, the preclearing reagent was removed, specimens were rinsed briefly with dH2O and then washed with PBSPC (phosphate buffered saline with added biocide ProClin300 (PBSPC, Sigma-Aldrich 48912-U) at 0,05% v/v) four times for 1.5 h, once overnight and again twice for 1.5 h at RT before proceeding to dehydration. At this step, samples could be stored in PBSPC at 4°C in the dark for up to four weeks without significant loss of fluorescence signals. Preclearing of non-perfused organs improved imaging for all tissues and was indispensable for the following organs (incubation time/notes): Spleen (4d), kidneys (4d), liver (4d), heart (4d, coronal section exposing all four chambers of the heart using a scalpel), large/blood-rich tumors (4d, larger than 500 µm in diameter), tongue (4d), lungs (2d), thymus (2d).

### 3D-Swiss Rolls for specimens of the GI tract

For specimens of the small and large intestine, the GIT was removed as a whole from the abdominal cavity by cutting the distal esophagus (approx. 3-5 mm from the stomach) and the rectum. Attached mesentery was removed by careful pulling with forceps or cutting. If the pancreas should be analyzed as a whole, the GIT was removed together with the entire pancreas and the spleen and then separated ex situ as a whole. Subsequent intestinal incisions were made at the pyloric sphincter, the ileocecal valve and distal of the cecocolic orifice, thereby separating stomach, small intestine, caecum and colon. Stomach and caecum were transferred to ice-cold PBSPC to slow down autolytic processes while the colon and small intestine were processed.

The distal end of the small intestine was gently pulled onto a rodent oral feeding gavage with ball-tip, attached to a 50 ml syringe containing ice-cold PBSPC. Holding the sample firmly on the gavage with fingers, chyme and feces were flushed out with ice-cold PBSPC. While flushing, the specimen was gradually and gently pulled onto the gavage to allow for thorough removal of feces also from the proximal end. After rinsing, small intestines were immediately flushed again and filled with NBF using a separate syringe with a feeding gavage. The small intestines were then laid out flat, forming an ‘N’, and cut into three equally long sections (SI 1 3). The created segments were transferred to a beaker containing NBF, noting the correct order of segments as well as the proximal and distal ends. The colon was processed as one single specimen.

3D-Swiss Rolls were formed as quickly as possible to prevent specimens from becoming too rigid for proper rolling. Rolling of the colon was conducted first, as it became too rigid for rolling shortly after immersion in NBF. To create 3D-Swiss Rolls, the specimens were cut open longitudinally along the mesenteric line and then transferred back into a flat dish containing NBF. Using forceps, the specimens were gently pulled over a wooden tampone swab (LP Italiana, 112298, cotton ends removed) with the luminal side facing outward and starting with the proximal end of the colon. Once rolled, the end of the swab was placed into the lower corner of a large sample-processing cassette (Macrosette M512, Simport) and the protruding end cut off at the opposite corner of the cassette.

This way, specimens could be placed diagonal in the cassette without touching the surface of the cassette (thus avoiding imprints in the tissue after fixation). The cassette was immediately transferred to NBF for fixation. Each section of the SI was processed as described for the colon, except rolling was started at the distal end of each segment to avoid excessive squeezing of the longer proximal villi. Stomach and caecum were then cleaned by inserting a gavage needle attached to a syringe containing ice-cold PBSPC into the stomach through the pylorus or caecum through the ileocecal valve, respectively. Contents were flushed out until organs were empty and rinsing buffer was clear. The specimens were filled with NBF and transferred into a flat histological cassette with a paper inlay for fixation in their physiological shape. In general, complete removal of chyme or feces from all the specimens is important because the plant-based nutrition of mice shows very high autofluorescence in LSFM imaging. At the same time, quick processing is even more essential during dissection to prevent autolytic damage of the tissues. Therefore, if not all residues of chyme or feces could be removed during dissection, further cleaning could be conducted after fixation.

After fixation, 3D-Swiss roll samples were unraveled from wooden holding sticks in a large bowl containing ice-cold PBSPC and remaining chyme and feces were carefully removed. Specimens were then rolled up again in the same orientation, now on plastic stirring spatulas (Brand, VWR 441-0217). This was necessary because the wooden cotton swabs used during dissection left dark marks on the tissues upon dehydration. Rolled specimens were transferred back to cassettes for dehydration and clearing. Specimens were then dehydrated following the quickBABB procedure described below. After dehydration, plastic stirring rods were removed before immersion in BABB because polystyrene does not withstand organic solvents.

### Processing of whole mice

For clearing of entire mouse bodies, animals were euthanized by CO_2_ inhalation and immediately perfused transcardially with 40 ml PBS with 10 IU Heparin (B. Braun, 25.000 IE/ 5 ml Heparin sodium, Melsungen, Germany) at 2 ml/min, immediately followed by perfusion with 60 ml NBF with 10 IU Heparin at 1.5 ml/min and again 20 ml PBS with 10 IU Heparin sodium at 2 ml/min to remove NBF. Mice were then decalcified by constant perfusion and immersion in 20% EDTA solution (Entkalker Soft, Carl Roth 6484.2) for six days at 2 ml/min in the dark. EDTA was removed by rinsing and perfusion with dH2O for 30 min. The skin was removed and the GI tract cleaned by incising at multiple locations and rinsing out contents with dH2O using a syringe with an oral feeding gavage. Mice were then immersed in the preclearing reagent at 30°C for 14 days with gentle agitation and exchanges of the reagent after three, six and nine days. To control evaporation, incubation was carried out in a container with airtight lid. After the last step, the preclearing reagent was discarded and animals briefly washed in PBSPC to remove bulk residues of the reagent. Mice were then washed in PBSPC for a total of 24 h: 3 x 3 h, overnight and again 2 h before proceeding to methanol (MeOH) based dehydration, delipidation and RI matching.

### Dehydration and refractive index (RI) matching (clearing)

Specimens were dehydrated using two different procedures, depending on the tissue. All individual organs except the brain were dehydrated using EtOH, Carl Roth, Cat. 0911) an automated tissue processor (Tissue-Tek VIP® 6 AI Vacuum Infiltration Processor, Sakura) inside histology cassettes. The custom protocol comprised of eight steps of 30 minutes each in a low-pressure environment to enhance diffusion of an increasing concentration series of EtOH: 70 %, 70 %, 80 %, 80 %, 90 %, 90 %, 100 %, 100 % (v/v). After dehydration, cassettes were dried using a paper cloth before RI matching by immersing specimens in BABB (one part benzyl alcohol (BA, Sigma Aldrich 305197-2L) and two parts benzyl benzoate (BB, Merck Millipore 8187011000)). Specimens were incubated in BABB in the dark for 24 h until fully transparent (less for very small or permeable tissues, two days for whole mice). Once cleared, specimens could be stored light-protected for at least three months at 4°C without loss of fluorescence signals.

Whole brains and whole mice were dehydrated manually at room temperature using methanol (MeOH, Merck Millipore 1060092511) and additionally delipidated with dichloremethane (DCM, Merck Millipore 1006681000). Brains were dehydrated at room temperature (RT) with gentle agitation in the dark at 20%, 40%, 60%, 80% and 100% methanol (v/v, diluted with dH2O, 1.5 h each) and again in fresh 100% methanol overnight at 4°C. Delipidation was carried out in 66% (v/v) DCM and 33% MeOH for 5 h at RT with gentle agitation and specimens briefly washed in 100% DCM for 15 minutes before immersion in BABB until fully cleared.

Whole mice were processed according to the same protocol but with longer incubation times of 4 h for the first two incubations (20 %, 40 % methanol), o/n (60 % methanol), 8 h (80 %), o/n (2 x 100% methanol and DCM/methanol) and 30 minutes (100% DCM). As higher concentrations of methanol evaporated more quickly, specimens were incubated in airtight glass containers.

### Light sheet fluorescence microscopy (LSFM), data conversion and visualization, scoring

Imaging was conducted using either a light sheet fluorescence microscope (LSFM) Ultramicroscope II® (UM2, LaVision Biotec, Bielefeld, Germany; now part of Miltenyi Biotec, Bergisch Gladbach, Germany) or LSFM Ultramicroscope Blaze® (UM Blaze, Miltenyi Biotec, Bergisch Gladbach, Germany). The UM2 instrument was modified compared to the original model at the excitation light path. Instead of six light sheets for excitation of fluorescence, the illumination light was channeled in only two light sheets that were oriented opposite towards each other. LSFM data was acquired using the UM2 for figures 1, 2, 4, 5, 7 and supplementary figures S2, S3, S6-9, S11-18. LSFM data was acquired using the UM Blaze for figures 3, 6 and supplementary figures S5 and S10. As light source, a supercontinuum white light laser SUPERK extreme EXR-20 (NKT Photonics) with a maximal power output of 2 W was applied combined with optical bandpass filters for fluorescence excitation and emission detection. For acquisition of anatomy using the UM2 tissues were imaged by excitation at 545 nm with a filter bandwidth of 20 nm and emission was detected at 595 nm with a filter bandwidth of 40 nm (545(20)nm → 595(40)nm), the anti-EpCAM IgG2a conjugated with AlexaFluor®750 was detected at 747(33)nm → 786(22)nm. Using the UM Blaze, anatomy was acquired at 520(40)nm → 572(23)nm and the antibody was detected at 740(40)nm → 824(55)nm. For imaging of entire mice, mosaic scans were conducted using the built-in feature of the UM Blaze and subsequent image stitching using the Imaris Stitcher version 9.3.1 (Oxford Instruments, United Kingdom). Raw image data in the .tiff file format was converted to the native Imaris file format using the Imaris file converter version 9.3 or higher and visualizations were created using Imaris version 9.5 or higher (Oxford Instruments, United Kingdom). Scoring of binding was conducted semi-quantitively by comparing maximum fluorescence signal intensities of each tissue in whole mouse scans of three mice.

**Figure 1.**
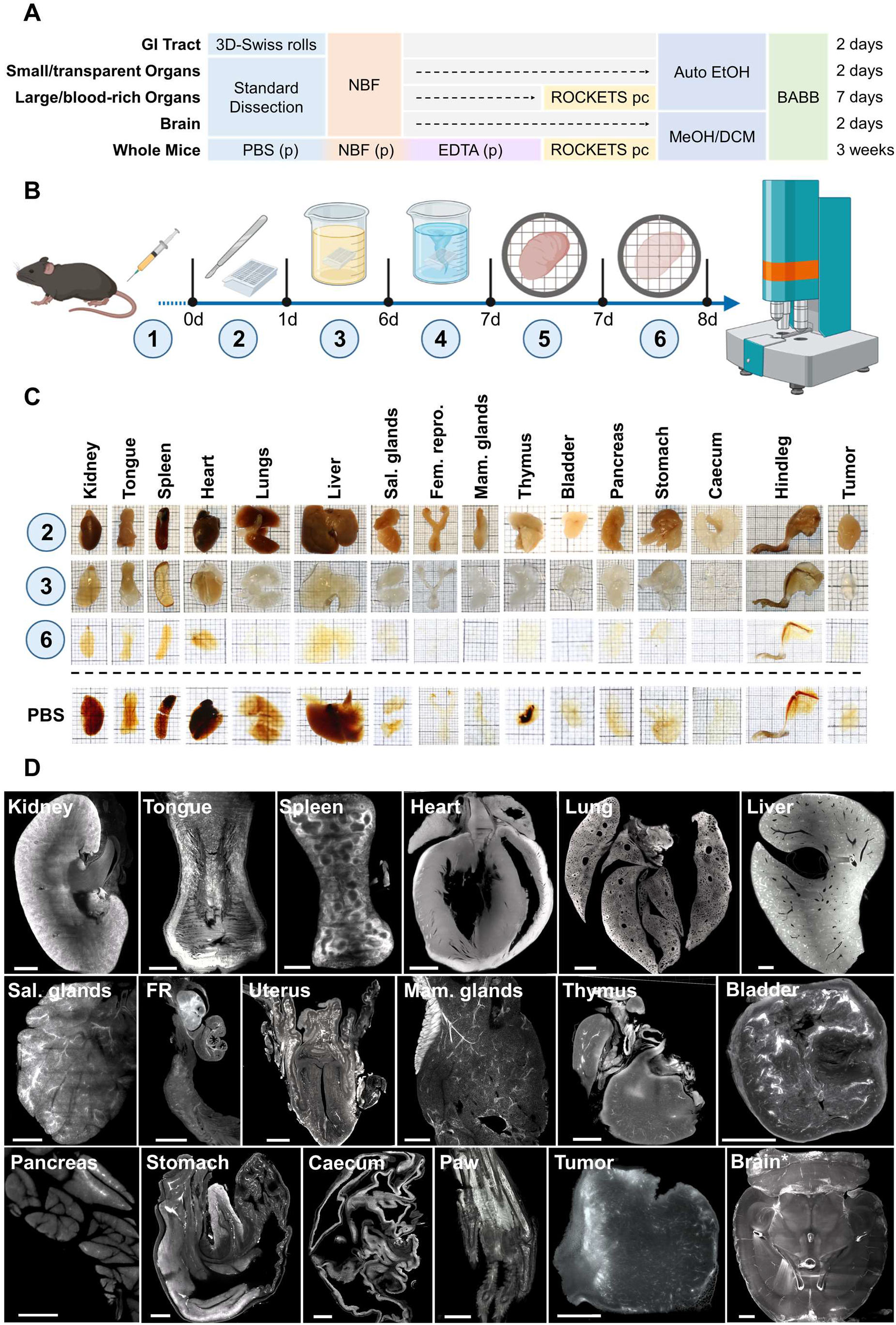
Modular clearing approach of the ROCKETS processing toolbox allows for simplified sample preparation for LSFM imaging. **(A)** Overview of presented procedures for processing and simplified clearing of mouse tissues or whole mouse bodies. GI tracts are processed using the 3D-Swiss Rolls procedure prior to fixation to enable holistic imaging. Other internal organs and tissues can be processed according to size and blood-content. Non-perfused large and blood-rich tissues are precleared using the developed preclearing reagent before dehydration (Fig. 1 B). Smaller tissues with less blood content do not require preclearing. All tissues except for the brain and whole mice are dehydrated with ethanol using an automated vacuum tissue processor (Auto EtOH). Due to its high lipid content, the brain is dehydrated in methanol and additionally delipidated using dichloromethane (MeOH/DCM). Only whole mice require perfusion to ensure timely fixation and decalcification of bones before the preclearing step. All specimens are cleared (RI matching) and imaged in BABB. Indicated times are total processing times from the day of dissection to cleared specimens. (p) = perfusion. **(B)** Workflow of passive preclearing of non-perfused murine tissues. Fluorescence labeled molecules are applied in vivo (1) prior to euthanasia, tissue dissection and fixation overnight (2). Fixed specimens are incubated in the ROCKETS reagent (3) and washed with PBSPC (4) before transfer to vacuum-enhanced dehydration (5) and RI matching with BABB (6). **(C)** Photographs of mouse tissues at indicated step of preclearing. Specimens are opaque and still contain blood pigments after fixation (2). After preclearing (3) samples are fully decolorized and swollen and become completely transparent after dehydration and RI matching (6). The bottom row shows tissues after dehydration and RI matching without preclearing (immersed in PBS). Particularly blood-rich organs are insufficiently cleared without perfusion or preclearing. Thick squares of the grid = 5 mm. **(D)** Maximum intensity projections (MIPs) of LSFM images (z = 50 µm) of the tissue’s autofluorescence (545 nm → 595 nm) at the widest diameter of precleared tissues. FR = Female reproductive organs (oviduct and ovary), Sal. glands = Salivary glands. All tissue areas could be imaged entirely without blurring. Scale bars = 1 mm.

### Processing of cleared tissue specimens for conventional 2D histology and slide scanning

Histological assessment after 3D-LSFM imaging was conducted by removal of BABB and washing with xylene (Roth, Cat. 9713.3) for 10 minutes before paraffinization using a Tissue-Tek VIP 6 Vacuum Infiltration Processor (Sakura). Paraffinized specimens were cut to 2.5 µm thick slices using a rotary microtome HM355S heavy duty with section transfer system and Cool-Cut module (Thermo Scientific). Floating tissue slices were picked up on glass slides and dried at 30°C o/n. Tissue sections were then deparaffinized and rehydrated automatically using a BenchMark ULTRA autostainer (Roche Ventana) programmed to apply: 3 x xylene (3 min.), 2 x 100 % ethanol (2 min.), 95 % ethanol (1 min), 70 % ethanol and dH2O (1 min). Hematoxylin and Eosin (H&E) stainings were performed using a VENTANA HE 600® system (Roche). The slides were imaged using slide scanner AxioScan® 7 (Carl Zeiss, Jena, Germany) and files were exported and visualized using ZEN Blue Edition Software, version 2.3 (Zeiss).

## 3 Results

### ROCKETS toolbox for passive clearing of mouse tissues

To develop a simple and harmonized protocol for large-scale and high-throughput LSFM for immunological research and preclinical drug development of all mouse organs or whole bodies for LSFM imaging we integrated, adapted, and complemented existing procedures for various tissues. To this end we focused on passive clearing techniques, which are often categorized by hydrophilic and organic solvent-based approaches (66–69) We chose to combine these concepts in a coherent two-step procedure, which we subsequently termed Rapid Optical Clearing Kit for Enhanced Tissue Scanning (ROCKETS). First, hydrophilic expansion (hyperhydration), delipidation and decolorization similar to the previously published CUBIC by Susaki et al. in 2014 (70) and, second, dehydration and organic solvent-based RI matching as described already in 1914 by Werner Spalteholz (71), which was first applied for modern LSFM of biological tissues by Hans-Ulrich Dodt et al. in 2007 (40) and later refined in the DISCO-family of clearing protocols, initiated by the work of Ali Ertürk et al. in 2012 (72–74). Subsequently, our developed ROCKETS toolbox allowed for choosing to process particular or all tissues of interest or even whole mice (**Fig. 1A**), for which each critical step is outlined below.

### Fixation

Tissue fixation in general is an important factor in tissue processing that has rarely been considered for tissue clearing. After *in vivo* i.v. administration of fluorescently labeled antibodies and euthanizing mice we fixed tissues using neutral buffered formalin (NBF), which covalently cross-links proteins (75) to keep bound antibodies linked to their target. We observed that the duration and temperature of fixation in NBF had a significant impact on clearing performance and undesired autofluorescence. Over-fixation (>12 h at room temperature (RT) or >24 h at 4°C) led to insufficient clearing, particularly of large and blood-rich tissues as well increased background fluorescence. Fixation <8 h at RT or <12 h at 4°C for large organs resulted in tissue damage during subsequent processing steps and lower specific fluorescence signal intensities in affected tissue regions. Thus, whole organs were fixed in NBF overnight at 4°C for 14-18 h immediately after dissection.

### Passive preclearing for large and blood-rich mouse organs

The goal of any tissue clearing protocol is to maximize transparency through reducing light absorbance and scattering (58, 68). The major source of absorbance in most biological tissues is hemoglobin, the pigment of red blood cells (66). Therefore, to flush blood from the vessels most clearing protocols start with transcardial perfusion of mice, a laborious and messy procedure (76, 77). To enable passive clearing and omit perfusion, we developed the concept of a hydrophilic preclearing step prior to dehydration and organic solvent-based RI matching. We reasoned that the original CUBIC cocktail as published by Susaki and colleagues in 2014 (70) and particular the aminoalcoholic component quadrol (*N,N,N′,N′-Tetrakis(2-Hydroxypropyl)ethylenediamine*) should be principally suitable to omit perfusion. The decolorizing ability of quadrol is based on releasing the light-absorbing prosthetic heme from erythrocytes (68). We found that the decolorizing effect for fixed whole liver lobes treated with various dilutions of quadrol generally increased with increasing concentrations, peaking at approximately 20 % quadrol (v/v, in dH_2_O) above which we observed no further improvement in effect nor time. Starting from the decolorizing reagent we rationally added further components to the mixture to reduce scattering and increase permeability. The cause for light scattering in biological tissues are inhomogeneous refractive indices, particularly between aqueous compartments, proteins, lipids and fatty acids (58). To elute different types of fats (delipidation) we added two surfactants, Tween-80® (T-80) and Triethanolamine (TEA) at 10 % (v/v), thereby avoiding commonly used octylphenol ethoxylates like Triton^TM^ X-100, which have been banned from using in the European Union by the European Chemicals Agency (ECHA) due to environmental toxicity. We further included urea as applied in the CUBIC reagent, which induces hyperhydration and corresponding swelling of the tissues, thereby increasing molecular flux and facilitating diffusion of all components through the tissues (68). As described previously for brain tissue (45) we also observed increased swelling of all organs with higher urea concentrations (not shown). At concentrations above 15% urea (w/v) we observed macroscopic deformations of large organs such as the liver (**Suppl. Fig. S1**). These morphological changes were permanent and not reversed through dehydration (and resulting shrinkage) and clearing. Therefore, we added urea at 10% (w/v), which was sufficient to induce reversible swelling without affecting anatomy. Dissolving of all components in deionized water (dH_2_O) yielded a highly viscous solution. To reduce viscosity, we incubated specimens at 30 °C and added 10 % dimethylsulfoxide (DMSO), which is known to promote both hydrophilic and lipophilic permeation through tissues (78). The final cocktail, which we termed *preclearing reagent*, was a yellowish solution with water-like viscosity at 30°C. Non-perfused mouse organs were incubated in 15 ml preclearing reagent per whole organ for two to four days at 30°C, depending on the organ. After treatment, all tissues except bone marrow appeared completely colorless, swollen and partially transparent (**Fig. 1C, step 3**). We detected splenic melanosis in some specimens, presenting as dark spots at one end of the spleen, as is frequently observed in mice with dark coat color (79) and which could not be removed through preclearing.

After washing, specimens appeared with a yellow-whitish non-transparent color and had re-gained their physiological size. After preclearing, we dehydrated and cleared specimens using a 1:2 (v/v) mixture of benzyl alcohol and benzyl benzoate (BABB, see below). After RI matching, all organs were fully transparent when treated using the preclearing reagent (**Fig. 1C, step 6**) and anatomy was unaffected (**Fig. 1C, step 6**; **Fig. 1D**). In LSFM scans of the autofluorescence at 545 → 595 nm precleared tissues could be imaged at high resolution throughout their entire volume (**Fig. 1D**) and showed overall higher fluorescence intensities compared to non-treated samples. Without preclearing, particularly large organs still contained significant amounts of blood and appeared generally more opaque after dehydration and RI matching (**Fig. 1C, PBS**). Without the introduced preclearing steps, several organs (kidneys, tongue, spleen, heart, lungs, liver, thymus, hindleg) could not be imaged entirely, particularly at lower wavelengths due to light attenuation and blurring towards the center (not shown). However, smaller organs or tissues with lower vascularization such as caecum, stomach, female reproductive tract, bladder, and lymph nodes were sufficiently transparent without preclearing. Thus, we concluded that preclearing of these organs could be omitted if imaging at higher wavelengths is intended (**Suppl. Tab. S1**). Yet, importantly, preclearing also improved image quality and signal-to-noise ratio (SNR) for small organs.

### 3D-Swiss Rolls for holistic assessment of the gastrointestinal tract

The sheer size and the convoluted tubular structure of the GIT, particularly the small and large intestine, makes it difficult to investigate microscopically. The GIT is neither structurally nor functionally a homogeneous tissue and it is therefore important to analyze it as a whole (80, 81). Therefore, we adapted the histological preparation technique of *swiss rolling* (82–84)) for LSFM-based three-dimensional imaging of the GIT. In reference to this, we termed the samples created by our technique *3D-Swiss Rolls*. After euthanizing mice, we removed the lower GIT as a whole and separated it *ex situ* into six specimens (**Fig. 2, steps 1, 2**): Stomach (STO), three equally long segments of the small intestine (SI 1-3), caecum (CAE) and colon (COL). Using an oral feeding gavage needle connected to a syringe, we flushed out chyme and feces and immediately filled the specimens with NBF (**Fig. 2, step 3**) to accelerate fixation and prevent autolytic processes. We used NBF instead of acidic Bouin’s fixative (as applied in the original procedures) to avoid fluorescence quenching and to streamline the workflow with processing of other organs. Next, we cut open the small intestine and the colon longitudinally (**Fig. 2, step 4a**) and rolled up the segments on wooden sticks with the luminal side facing outward and further fixed them in this position (**Fig. 2**, **steps 4b and 5**). Quick processing turned out as essential during all steps of the procedure. If not processed quickly, particularly the stomach and the proximal third of the small intestine started to deteriorate within minutes due to autolytic processes from exposure to gastric acid, bile and digestive enzymes as observed previously (85). Otherwise, resulting damages to the tissues’ microanatomy due to slow tissue processing might be misinterpreted for toxicity-related effects of investigated drugs. Also, 3D-Swiss Rolls had to be placed carefully into histology cassettes without being pressed against the surface to avoid imprints on the specimens (**Suppl. Fig. S2**). Once fixed, the rolls could be handled with less caution and retained their rolled form during washing, change of holding sticks and automated dehydration. After dehydration and RI matching, samples were stiff and could be easily mounted for LSFM imaging. We confirmed the anatomical integrity of the GIT specimens by LSFM imaging as well as slide-based histology with hematoxylin and eosin (H&E) staining (**Suppl. Fig. S3A and S4**). High-resolution LSFM of 3D-Swiss Rolls allowed us to identify individual cells (enterocytes, goblet cells and paneth cells) and single nuclei in the entire GIT without additional counterstaining (**Suppl. Fig. S3A, C**).

**Figure 2.**
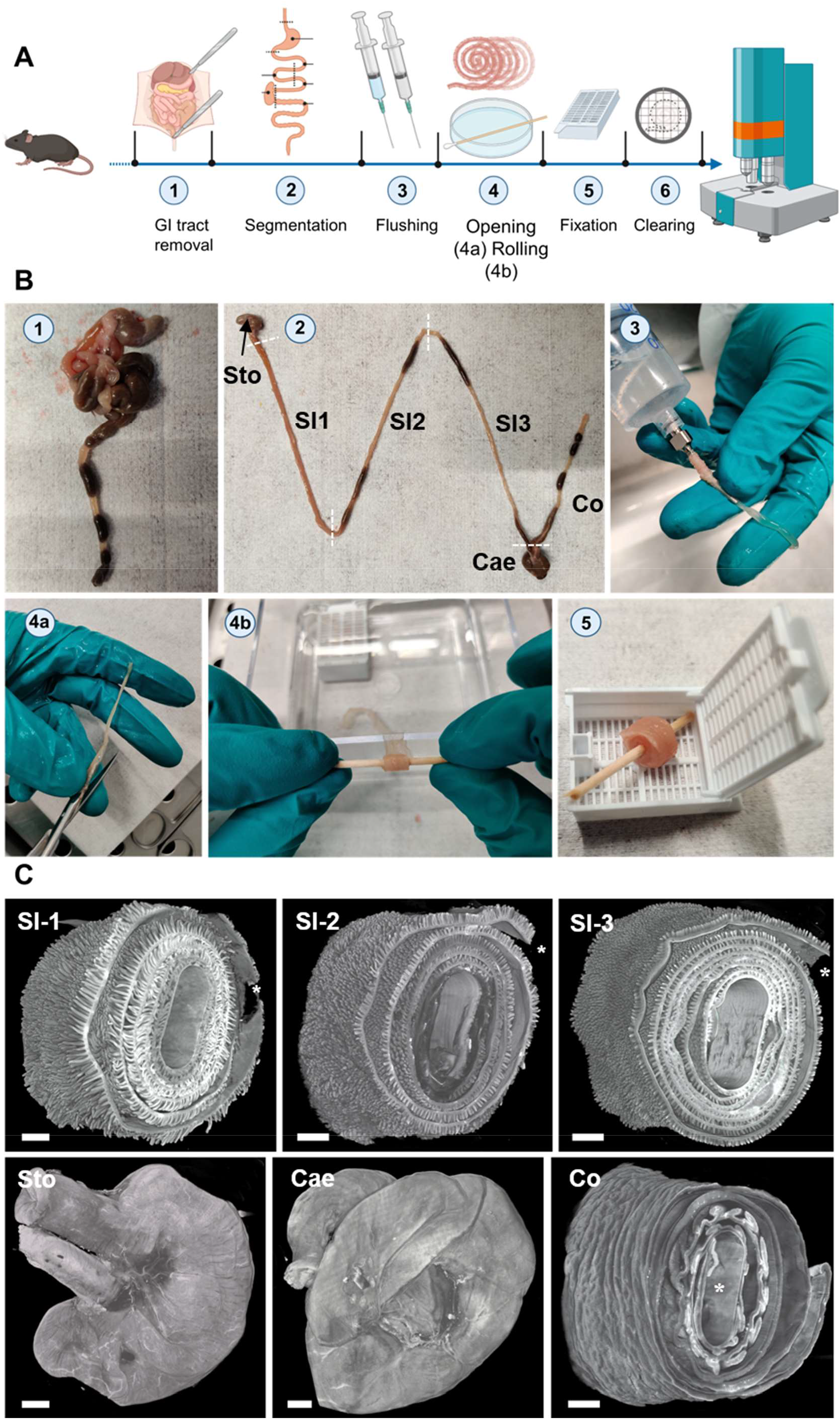
3D-Swiss Rolls sample preparation procedure for LSFM imaging enable holistic assessment of the entire GIT. **(A)** Schematic and **(B)** photographic representation of the 3D-Swiss Rolls workflow. (1) After euthanasia the lower GIT is disconnected from the body by incisions at the esophagus and rectum and removed entirely. (2) Six specimens are created by cutting as indicated by dashed lines: stomach (STO), three segments of the small intestine (SI 1-3), caecum (CAE) and colon (COL). (3) Each specimen is cleaned by flushing out chyme and feces with PBS_PC_ and then immediately filled with NBF for fixation. (4a) SI and COL segments are cut open along the mesenteric line and (4b) rolled up on wooden sticks to create 3D-Swiss Rolls. (5) The created 3D-Swiss Rolls are then fixed without touching the surfaces of the histology cassette for 14-18 h in NBF at 4°C. After fixation, 3D-Swiss Rolls are unwound and re-rolled on plastic stirring rods for dehydration and clearing (not shown). **(C)** Surface rendering of LSFM image stacks of the tissue autofluorescence (545 → 595 nm, grey). 3D-Swiss Roll segments of the small intestine (SI1-3) and colon (Col). Stomach (Sto) and caecum (Cae) retained their physiological form. *proximal end of the organ in 3D-Swiss Rolls. Scale bars = 1 mm.

### Clearing of whole mouse bodies

Next, we asked whether we could also apply the ROCKETS procedure even to whole body LSFM. In this case, we reasoned to first perfuse mice to avoid autolytic processes and to ensure rapid tissue fixation. Therefore, in contrast to the perfusion-free whole organ-clearing protocol, we perfused whole mice with NBF to ensure timely and thorough fixation, followed by 25% ethylenediaminetetraacetic acid (EDTA) to elute light absorbing calcified minerals from bones, similar to previous reports (86–88). Subsequently we removed the skin and cleaned the GI tract from chyme and feces *in situ* before mice were incubated in the preclearing cocktail for 10-14 days with three exchanges, using a sealable container to prevent excessive evaporation. After washing off the preclearing reagent, we dehydrated and delipidated the mice before RI matching with BABB as described in the next paragraph. The procedure resulted in excellent transparency of entire mouse bodies (**Fig. 3A**) and all inner organs could be easily identified in LSFM scans (**Fig. 3B**).

**Figure 3.**
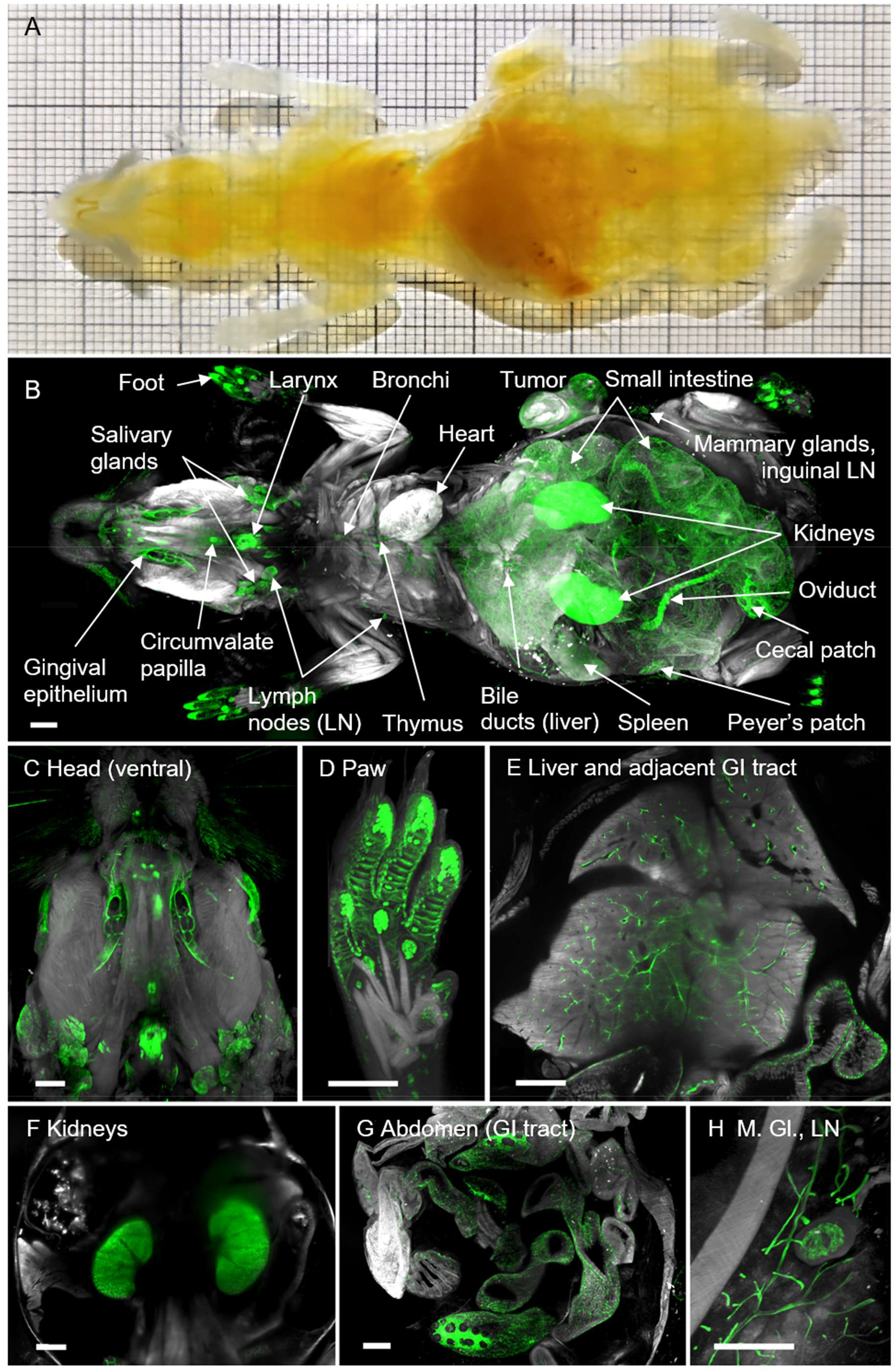
Whole mouse body cleared using the ROCKETS whole-mouse procedure and LSFM imaging reveals holistic biodistribution of anti-EpCAM antibody (G8.8R). **(A)** Mouse (ventral view) body after decalcification, preclearing, dehydration and immersion in BABB shows excellent transparency. Thick squares of the grid = 1 cm. (**B**) LSFM rendering of the tissue’s autofluorescence (grey) and anti-EpCAM staining (G8.8R in green) as overlay enabled quick localization of antibody disposition and identification of positive tissues. **(C-H)** LSFM renderings (ventral views) of EpCAM^+^ tissues (G8.8R in green) *in situ*. M. Gl. = Mammary glands, LN = Lymph node. Scale bars = 2 mm.

### Dehydration, delipidation and RI matching

For dehydration of individual organs (with or without preclearing) we used a tissue processor, which automatically executed dehydration within 4.5 h enhanced by negative pressure (vacuum) and as described previously (48). Brains and whole mice required additional delipidation and, therefore, we manually dehydrated these tissues by adapting the previously published iDISCO+ protocol (89) using methanol (MeOH) and dichloromethane (DCM) (73, 90). Importantly, we omitted the previously described bleaching step with H_2_O_2_ of the original iDISCO+ protocol to avoid rapid quenching of fluorophores induced by oxidative treatments. Irrespective of the applied ROCKETS modules, all specimens were finally immersed in BABB for RI matching and imaging.

### Biodistribution of an anti-EpCAM antibody (G8.8R)

The cell surface glycoprotein EpCAM is highly expressed on a variety of epithelial cancers but also in healthy tissues, successful therapeutic targeting relies on balancing on- and off-tumor effects. To map EpCAM expression throughout the whole organism, we employed our newly established ROCKETS procedure to investigate the biodistribution of the monoclonal anti-EpCAM IgG2a antibody (clone G8.8R, conjugated with AlexaFluor750®) after i.v. application into the tail vein of wild type C57BL/6 mice bearing a subcutaneous ectopic tumor (EpCAM expressing pancreatic cancer cell line KPC-4662). Upon analysis of the biodistribution, we scored the detected binding levels based on fluorescence intensity levels (**Suppl. Tab. S2**). To account for inherent signal contribution of the autofluorescence, we always scanned negative controls of the same tissue (without antibody) that were equally processed according to the ROCKETS protocol. First, we created LSFM-based 3D-renderings of entire mice and mapped EpCAM-(G8.8R)-positive tissues throughout the body (**Fig. 3B-H**). Hereby, we could determine individual EpCAM^+^ organs and structures: oral cavity (gingival epithelium) and tongue (gustatory papillae), larynx, thymus, salivary glands, trachea, thymus, bronchi and bronchioles, pancreas, liver (bile canaliculi and gall bladder), gastrointestinal tract (stomach, small intestine, caecum, colon and rectum), kidneys and urinary tract, female reproductive organs (oviducts), mammary glands, foot pads (sweat glands), hair follicles, brain ventricles (choroid plexus) and tumor.

Based on our findings in intact mice, we processed and cleared whole organs individually according to the ROCKETS toolbox to investigate biodistribution of the EpCAM-(G8.8R)-antibody at higher magnifications on a cellular level. All tissues that we considered negative in whole mouse imaging also proved negative upon individual inspection (connective, muscular and nervous tissues, bones). Cuboidal and columnar epithelia clearly stained for EpCAM-(G8.8R), as well as lymphoid organs (thymus, lymph nodes (LNs), Peyer’s patches (PPs) and spleen)(**Fig. 4**). Thereby, all known EpCAM^+^ tissues in mice (91–94) were accessed and bound by the anti-EpCAM antibody clone G8.8R *in vivo* within 24h of circulation. We investigated binding in each organ in detail and could easily determine substructures and individual cells that were positive for the antibody **(Fig. 4-7, Suppl. Fig. S6-18)**. For example, we identified individual nephrons in the kidney and determined that binding was restricted to distal convoluted tubules and collecting ducts with distinct binding to intercalated cells and excluded from proximal convoluted tubules and glomeruli **(Suppl. Fig. S12)**. On a subcellular level, binding was restricted to basolateral membranes in all positive epithelia (**Suppl. Fig. S5**), which also reflects known EpCAM-expression patterns (30). Furthermore, we detected more pronounced binding to proliferative stem cells in crypts of the small intestine than at differentiated enterocytes and goblet cells in the villi, corresponding with described EpCAM downregulation upon differentiation in the GI tract (95). Similarly, binding levels gradually decreased from the bottom of the crypts in the caecum and colon towards the luminal surface (**Fig. 7C, D**), also corresponding with respectively reported EpCAM expression gradients in rats (96). In lymph nodes, spleen, thymus and inside PP follicles we detected non-polarized membranous and diffuse signals (**Suppl. Fig. S6-S8**) that may be attributed to low EpCAM expression on T-, B- and dendritic cells in mice (92–94) but may to certain degree also reflect Fc-dependent binding of the antibody. Within the tumor, EpCAM-binding appeared characteristic for carcinomas (30) as non-polarized and highly heterogeneous (**Suppl. Fig. S10)**.

**Figure 4.**
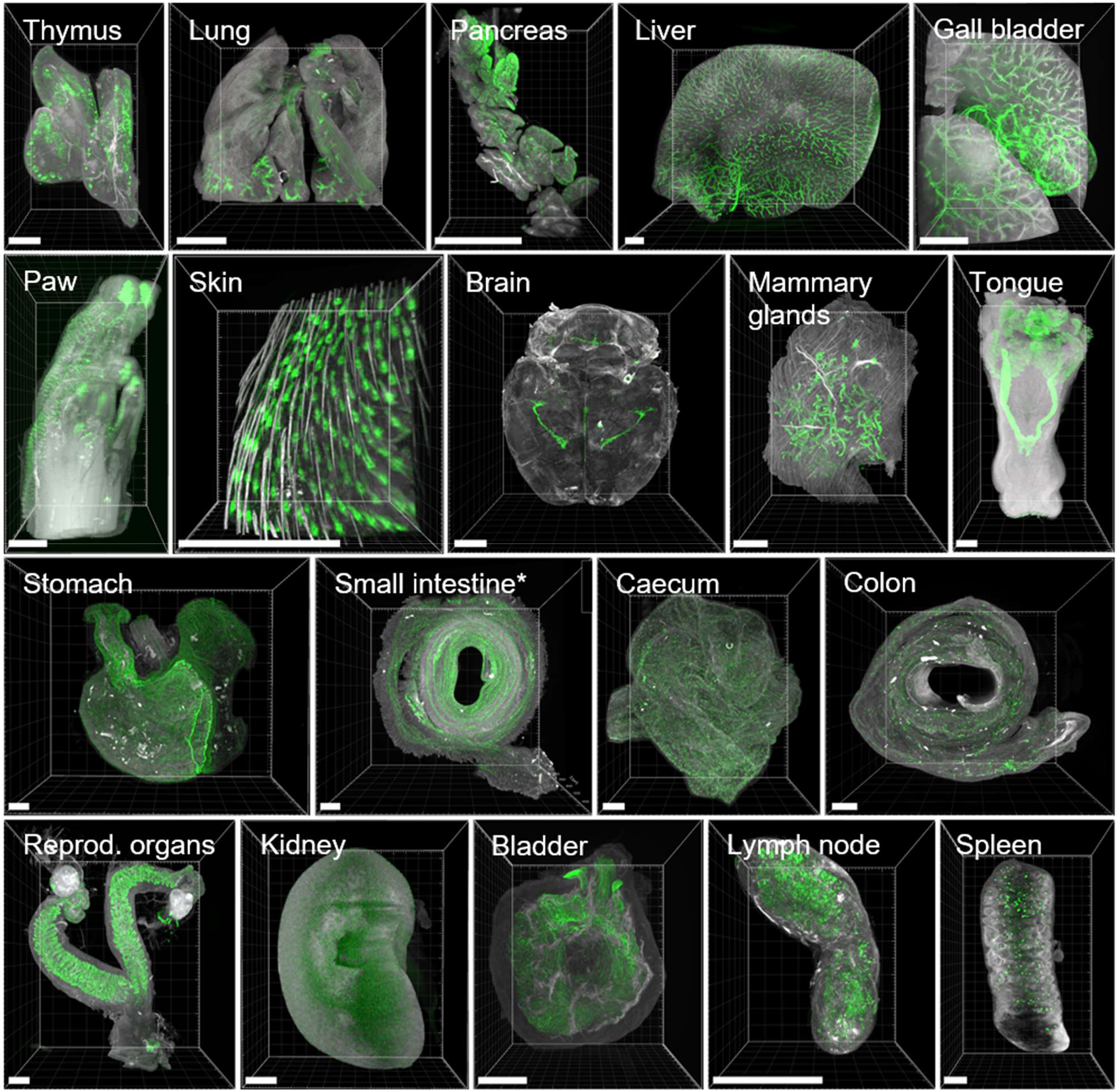
3D-Renderings of LSFM images show highly heterogeneous binding of anti-EpCAM antibody (G8.8R) between and within organs. EpCAM-stainings (G8.8R in green) and tissue anatomy revealed by tissue autofluorescence (grey) in maximum intensity projections (MIPs) of selected positive tissues. EpCAM-binding was detected at (but not limited to) previously published sites of EpCAM expression (91). *Only the first of three segments of the small intestine depicted (corresponding to duodenum and proximal jejunum). Scale bars = 1000 µm.

Of note, we further detected anti-EpCAM-(G8.8R)-stainings in tissues that previously had not been investigated for EpCAM or had even been reported negative. We observed EpCAM-expression at all types of gustatory papillae (fungiform, circumvallate and folate, **Fig. 5C, E**), which were not addressed in published expression analyses in both mice and humans (28, 97). Mucous salivary gland (MSG) acini were found EpCAM^+^ in some histological studies (98) while others did not detect EpCAM (99) or did not discriminate between mucous and serous salivary glands (SSG) (100). In LSFM scans of lingual salivary glands we observed a heterogeneous binding pattern across the entire gland and generally much lower signals in MSG than in SSG acini (**Fig. 5F-I**). Similarly, EpCAM expression in choroid plexus (CP) epithelia was not analyzed in investigations of the human brain (100) or had been even described as EpCAM-(98). However, we detected high levels of EpCAM-(G8.8R)-binding to individual CP cells (**Fig. 6**) distinctively delineating the CPs in mouse brains.

**Figure 5.**
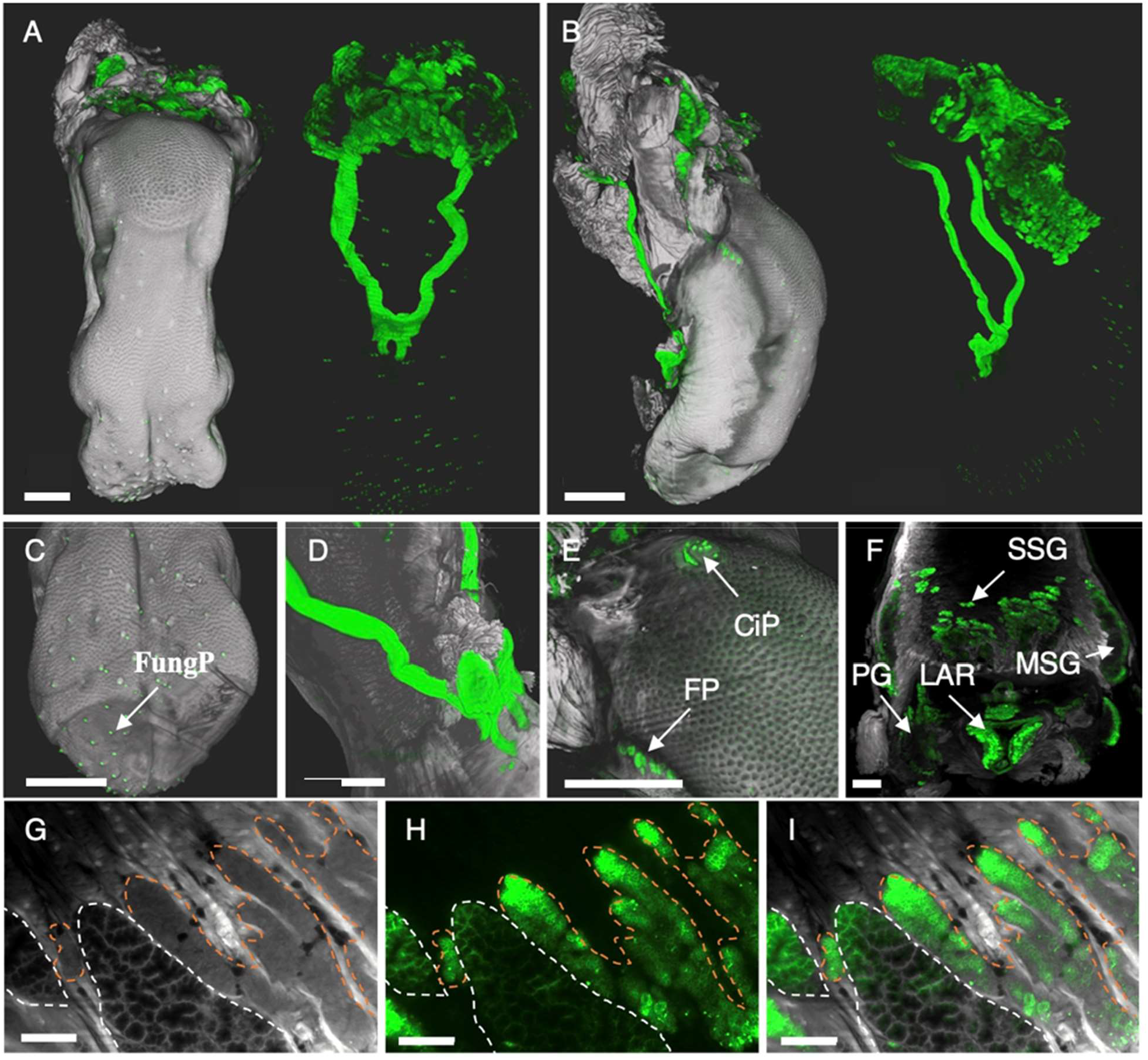
3D-Renderings and single LSFM images display highly heterogeneous binding of anti-EpCAM antibody G8.8R to the tongue and salivary glands. **(A)** Dorsal and **(B)** lateral view of surface renderings of the tongue and associated tissues. Left images depict renderings of the tissue anatomy (grey) and the bound anti-EpCAM antibody (G8.8R in green) as overlay. Right images depict only the antibody signal (green) without anatomical context. **(C)** Tip of the tongue with positive gustatory fungiform papillae (FungP) **(D)** Positive sublingual excretory ducts at the tongue bottom **(E)** Circumvallate papilla (CiP) and folate papillae (FP) **(F)** Mucous salivary glands (MSG) and serous salivary glands (SSG), parotid gland (PG) and larynx (LAR). **(G)** Single digital section of the tongue depicting both mucous (white dashed lines) and serous (orange dashed lines) salivary gland anatomy, **(H)** bound G8.8R (green) and **(I)** overlay of both channels. Scale bars = 1 mm (A-F) and 150 µm (G-I).

**Figure 6.**
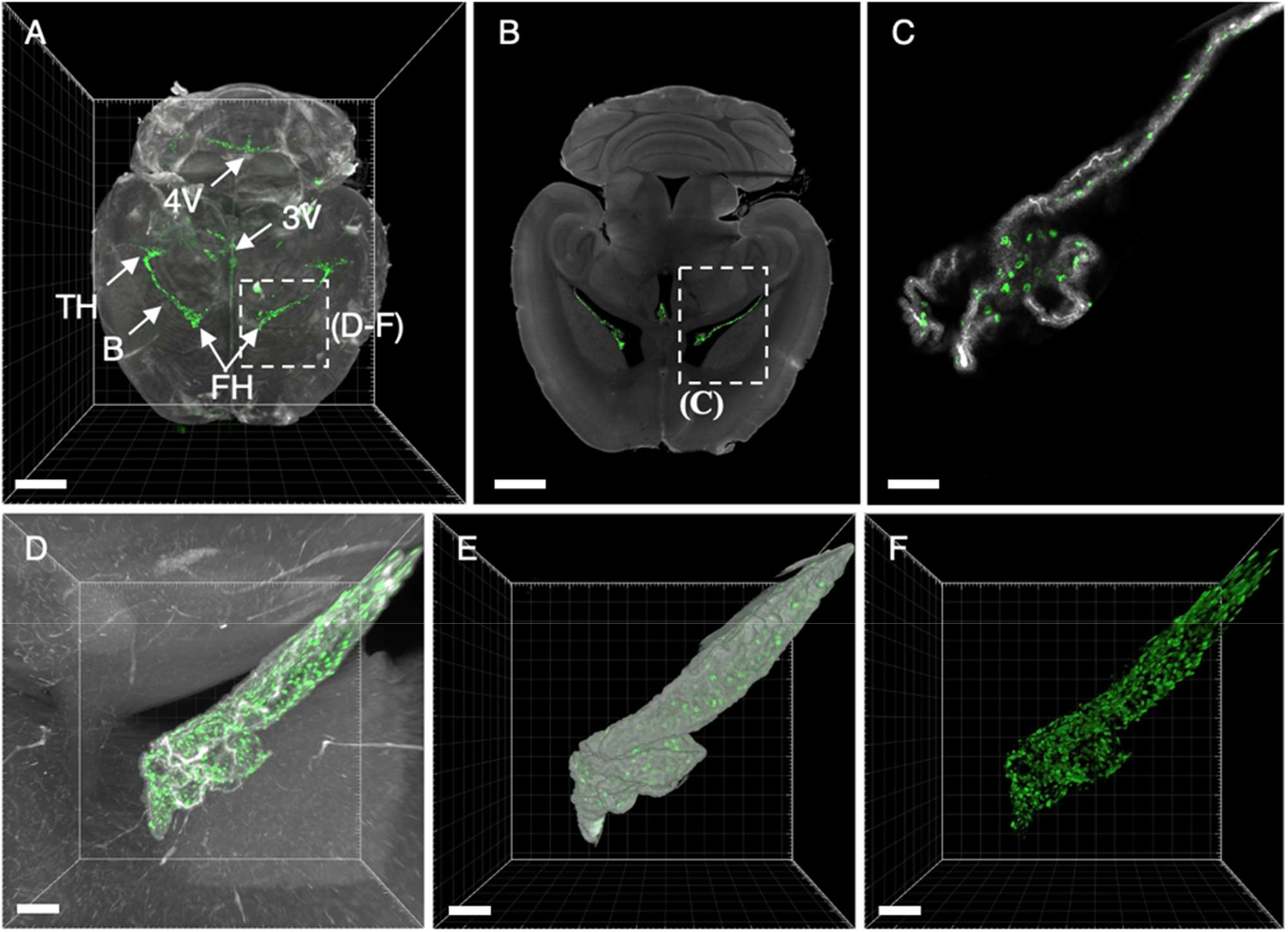
3D-Renderings and single digital sections reveal EpCAM-binding to choroid plexi in the brain. **(A)** Dorsal maximal intensity projection (MIP of the entire brain anatomy derived from the autofluorescence (grey) and binding of the EpCAM-specific antibody G8.8R (green). **(B)** Single LSFM image reveal anti-EpCAM antibody binding to choroid plexi of the temporal horn (TH), frontal horn (FH), 3rd ventricle (3V), 4th ventricle (4V, in A) and body (B, central part). **(C)** Higher magnification image of area indicated in image B display binding to individual choroid plexus cells. **(D)** Maximum intensity projections (MIP) and **(E, F)** surface renderings of the entire frontal horn choroid plexus with bound anti-EpCAM antibody (G8.8R) extending into the ventricular space as indicated in image (A). Scale bars = 2 mm (A, B) and 100 µm (C-F).

LSFM-scans of 3D-Swiss Rolls allowed us to holistically investigate binding in the entire GIT without the requirement of physical sectioning (**Fig. 7**). The stomach showed a highly heterogeneous EpCAM-binding pattern, pronounced at the glandular mucosa directly adjacent to the limiting ridge and in the gastric epithelium throughout the glandular stomach (**Fig. 7A and B**). The cornified, stratified squamous epithelium of the forestomach showed very weak signals. In the small intestine the binding patterns and levels were similar throughout the entire length and circumference, restricted to basolateral membranes of epithelial cells (**Fig. 7B**). In the small intestine the binding patterns and levels were similar throughout the entire length and circumference, restricted to basolateral membranes of epithelial cells and prominent in the crypts (**Fig. 7B**).

**Figure 7.**
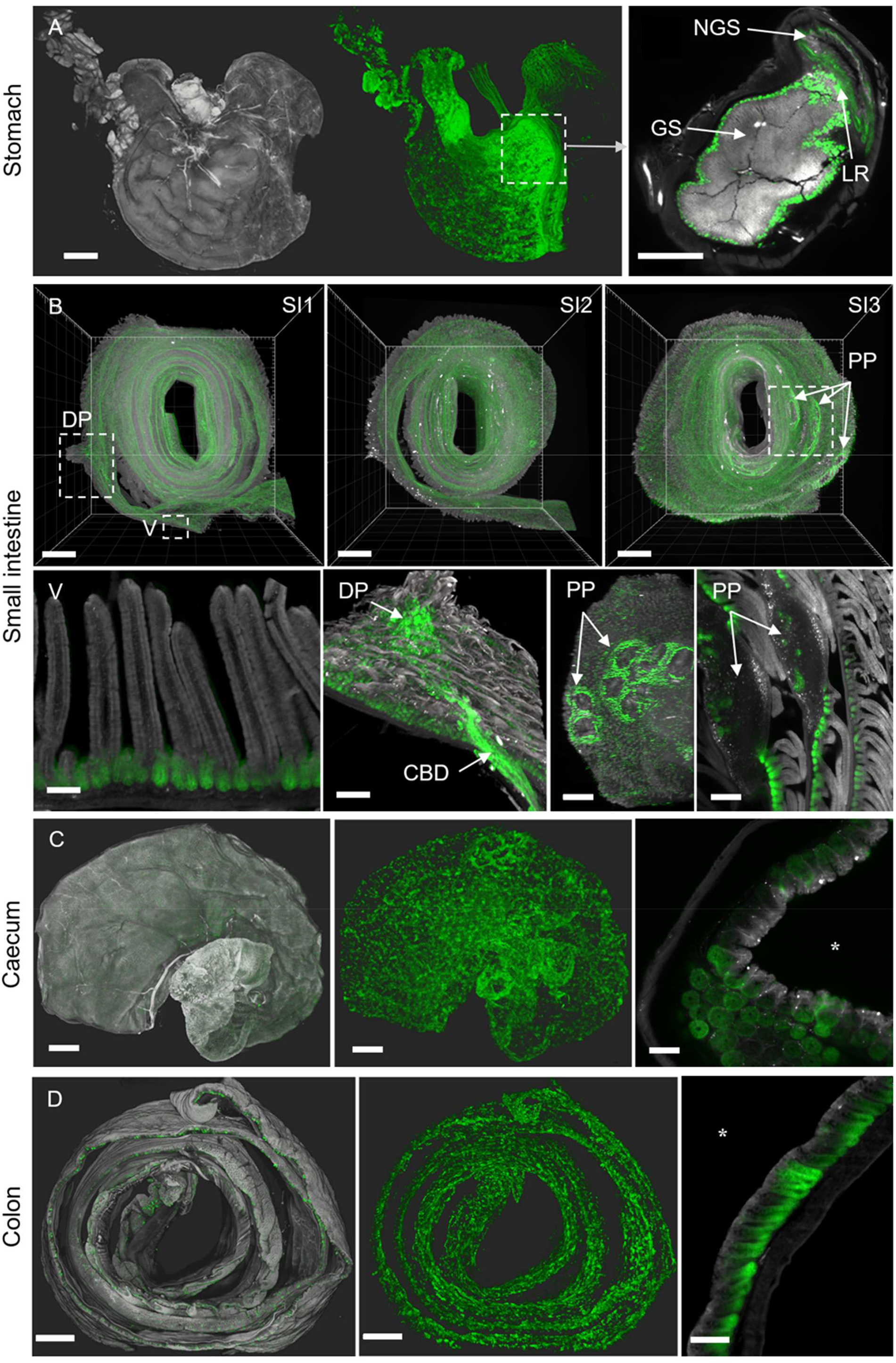
3D-Swiss Rolls present anti-EpCAM-staining (G8.8R) in the GIT. Tissue autofluorescence (grey) and anti-EpCAM-staining (G8.8R in green). **(A)** Surface renderings of the stomach and associated tissues. Left image depicts tissue anatomy, middle image depicts anti-EpCAM staining (G8.8R) antibody in the same specimen without anatomical context. Right image shows junction of the glandular (GS) and non-glandular stomach (NGS) with increased binding to the glandular mucosa near the limiting ridge (LR) **(B)** All three segments (SI1-3) of the small intestine as 3D-Swiss Rolls (upper row) with indicated duodenal papilla (DP) and several PP) exposing increased anti-EpCAM-binding. Lower row shows higher magnifications of structures as indicated in B. Binding was restricted to basolateral membranes of epithelial cells with pronounced binding to the crypts and decreased or no binding in the villi (V). Anti-EpCAM-binding was increased in crypts of the major duodenal papilla, common bile duct (CBD) and near PP compared to overall binding levels in the small intestine. Signals within PP follicles were diffuse and non-membranous, as observed for other lymphoid organs (Suppl. Fig. 6-9). **(C)** Caecum and **(D)** colon show similar patchy binding patterns with decreasing gradients from the proliferative bottom of the crypts towards the luminal surface of the tissues (arrow, luminal domain indicated by *asterisk). Scale bars = 1 mm (A, B SI1-3, C, D) and 100 µm (C-F).

However, in deviation from gross binding patterns, we detected strong and distinct EpCAM expression at the critical junction where the common bile duct and pancreatic duct drain into the small intestine, namely the major (papilla of Vater) and minor duodenal papillae, but also significantly increased EpCAM levels in the common bile duct (CBD), and in proximity of PPs (**Fig. 7B**, lower row). Interestingly, the PP dome epithelium showed low binding levels at the edges and was completely negative at the very center. Within PP follicles signals EpCAM-binding patterns appeared very similar to lymphoid follicles in LNs, corresponding to their shared immunological function (**Suppl. Fig. S6-S8**).

## 4 Discussion

In this work, we have integrated current knowledge and advanced procedures in tissue clearing to create a substantially simplified, streamlined and versatile sample preparation toolbox for LSFM that we termed ROCKETS. Experimenters may choose a suitable protocol from the ROCKETS toolbox for any mouse organ of interest or entire mouse bodies. The modular manner to apply the appropriate procedure and application should help to efficiently analyze any tissue type of interest or even all mouse organs in a standardized high-throughput mode relevant for basic immunological research but also for thoroughly assessing targets and reagents for novel theragnostic strategies in the preclinical development stage.

For assessing very large and blood-rich organs ROCKETS provides the advantage of efficient clearing with the developed passive two-step approach, which allows to omit transcardial perfusion, which is required for most other published protocols (69). Particularly for large-cohort preclinical animal studies, this simplification is an important element to reduce complexity and effort for tissue clearing. However, our chemical decolorization approach by eluting light-absorbing components does not necessitate perfusion only in terms of optical clearing. Blood remains inside the vessels, which has to be considered when fluorescence-labeled antibodies are applied via intravenous injections. In our case, the applied anti-EpCAM antibody was fully cleared from the bloodstream within 24 hours.

For smaller and less vascularized tissues the chemical preclearing step can be entirely omitted to reduce incubation times, waste and expenses. However, clearing of all tissues generally benefits from the preclearing through increased signal-to-noise ratios, which helps to enhance the measurement of even discrete specific signals. Of note, for direct comparison of different tissues within a given experiment, all samples should be treated equally to ensure comparability of fluorescence signal intensities. Apart from sample size and type, the choice of fluorescence probes generally affects clearing requirements. Red or near-infrared emitters may be detectable at high contrast while blue or green emitters can appear blurry because light at respective wavelengths interacts more with biological tissues and is therefore scattered (101). Furthermore, because of high autofluorescence levels of biological tissues in the blue-green color spectrum, signal-to-noise ratios might not allow for reliable detection in respective channels, which should be considered for each study. In our study, we chose AlexaFluor-750 as the fluorescence dye to label the biodistribution of an EpCAM-specific antibody, which emits in the near-infrared spectrum. Therefore, this reporter was well suited for sensitive detection deep within large organs and even in whole mice.

Histological investigations of the murine GIT are mostly performed using thin slices of tissue fragments or conventional *Swiss rolls* (82–84) or only focused on particular areas of the intestinal tract (52, 63) and thus, inherently underrepresent its three-dimensional complexity. 3D-Swiss Rolls allowed us to clear and image the small intestine and colon in full length and circumference at cellular resolution without affecting its microanatomy. The holistic imaging revealed that antibody binding was significantly elevated in the vicinity of functionally critical structures like Peyer’s patches and particularly at the duodenal papillae, which are difficult to locate on histological slices. However, the procedure required quick handling and processing to halt autolytic processes, which take place in gastrointestinal tissue specimens as a result of exposure to gastric acid, bile and digestive enzymes (85). Thus, any study using 3D-Swiss rolls should be well prepared and the technique practiced in advance, particularly because autolytic damage to the tissues may be mistaken for drug-induced lesions later on.

After dissection and optional preclearing, all organs except brains were dehydrated automatically in a tissue processor without user interference, which further streamlined and simplified the overall process (48). The brain had additionally to be delipidated using MeOH and DCM because of its lipid-rich composition. It should be noted that this difference in dehydration might affect comparability between the brain and other organs in terms of signal intensity. After dehydration, all specimens were cleared using BABB and could therefore be imaged without exchanges of the immersion medium during imaging.

As opposed to tissue processing for single organ imaging, whole mouse bodies required to ensure timely and thorough fixation as well as decalcification of bones. The remaining clearing process for whole mice was overall simple and fully passive and we could process multiple animals in parallel, limited only by the number of available perfusion pumps. *Ex vivo* LSFM imaging of cleared mouse bodies provided significantly higher resolution than typical *in vivo* imaging methods (102) but also produced very large data sets of several hundred gigabytes of data per animal. Correspondingly, data handling and three-dimensional rendering required significant computing power but then enabled holistic and highly detailed assessment of the biodistribution of the anti-EpCAM antibody for straight-forward identification of positive and negative tissues.

Biodistribution mapping showed that all known EpCAM^+^ tissues in mice (91–94) were specifically labeled with the EpCAM-antibody clone G8.8R 24h after in vivo administration. Imaging of entire animals also allowed for direct comparison of fluorescence intensity levels to derive semi-quantitative binding scores throughout the body. Accordingly, we observed significant differences in absolute intensity levels between animals but, importantly, the relative intensity distribution between body regions was equal for all investigated mice.

High-resolution imaging of cleared whole organs confirmed the findings in whole mice and provided more detailed information about binding patterns at a cellular level. Binding in all simple and pseudostratified epithelia was restricted to basolateral membranes, in accordance with known expression patterns (30, 96). The detected signal intensities corroborated published differences in cellular expression levels of EpCAM, which are generally higher on proliferating cells and gradually downregulated upon differentiation (95). This pattern was clearly observed across the small and large intestine, where EpCAM(G8.8R)-staining gradually subsided from proliferating zones at the bottom of the crypts towards more differentiated cells of the apical domain. These results underline the high sensitivity of LSFM and great potential for quantitative binding analyses in general.

Importantly, utilizing our ROCKETS procedure we uncovered in our comprehensive EpCAM-biodistribution studies, highly positive EpCAM tissue sites that have either not been sampled in published histological expression analysis or have been explicitly reported as EpCAM^−^ in mice or humans (29, 92, 94, 98, 100). All types of gustatory papillae, which represent clusters of specialized epithelial cells, known to express EpCAM in chickens (103, 104), were also EpCAM^+^ in mice and – considering the conserved expression in other epithelia – likely in humans.

For salivary glands, some expression analyses did not differentiate between types of salivary glands (100) or defined MSG as EpCAM^−^ (99). In LSFM scans, we detected significant differences in binding levels between lingual SSG (high) and directly adjacent MSG (negative or low). Therefore, we determine MSG as weakly EpCAM^+^ in mice, and attribute seemingly contradictory negative EpCAM stainings of MSGs in histological studies (99, 100) because of under-sampling or masking/loss of epitopes upon cross-linking fixation or processing.

In the brain, we detected no EpCAM expression in nervous tissue, in agreement with reports of human brain samples (100). However, we observed clearly EpCAM^+^ CP cells inside all ventricles in contrast to early reports of CP cells and ependymal cells as EpCAM^−^ (98). CPs comprise of simple cuboidal epithelium (105) and express various cell adhesion molecules that are generally associated with EpCAM in all other epithelia (e.g. E-Cadherin) (106). Furthermore, the blood–cerebrospinal fluid (CSF) barrier is implemented by tight junctions between CP cells (107), which are formed under contribution of EpCAM in all other tissues (108). In contrast to nervous tissue of the brain, CP cells can be considered accessible for antibodies because the CP vascularization comprises of fenestrated endothelium, which is generally leaky for macromolecules like the investigated antibody G8.8R (109). Therefore, we define CP cells as EpCAM^+^ in mice as visualized by LSFM imaging.

In summary, LSFM imaging provided unprecedented holistic insight into the biodistribution of an intravenously administered antibody. The great sensitivity and the readily discovered novel binding sites underscore the analytical power and broad spectrum of applications for LSFM imaging in drug discovery. As we discovered duodenal papillae as sites of high EpCAM expression, our results may have far-reaching implications for preclinical studies and clinical translation of EpCAM-targeted therapeutics as these are the sites digestive enzyme release from the pancreas. Many clinical studies targeting EpCAM did not reach their primary endpoints in the past due to dose limiting toxicities like pancreatitis (110, 111) or gastrointestinal-related adverse events (27, 112, 113). In light of our results, even neurotoxicity that in the past had been attributed to vascular-vascular-leak syndrome or presumed non-specific binding may deserve re-assessment considering the high level of EpCAM^+^ CP cells as important sites for cerebrospinal fluid secretion in the brain (114). Thus, future studies will require to particularly scrutinize immunotherapies whether they also target concomitantly these potential sensitive anatomical locations.

ROCKETS combined with LSFM imaging provides a highly versatile analytical platform for drug discovery. Generally, the described ROCKETS toolbox may be applied for preclinical assessment of any therapeutic compound or other fluorescence-labeled molecules. The methods are simple, make use of cheap reagents and provide sufficient throughput for large-scale studies. Importantly, the procedures are non-destructive for the investigated specimens. Therefore, LSFM imaging can be incorporated into existing preclinical analytical workflows. We envision ROCKETS and LSFM not to replace but rather complement gold-standard histological analyses.

However, the technology also carries some inherent limitations to be considered in each study. For example, intravenous administration of labeled antibodies is of limited use for actual expression studies because of potential inaccessibility of target cells *in vivo*. If target expression should be investigated, additional *ex vivo* immunofluorescence stainings of large tissue specimens may be conducted as described elsewhere (74, 115, 116). However, this approach bears limitations on its own because slow antibody diffusion into large tissue specimens still represents a major burden for *ex situ* staining. We did not investigate if the developed clearing methods preserve fluorescence signals from endogenous reporter proteins but it is likely that the organic solvent-based clearing would diminish fluorescence signals as described previously (69). In the future, further development may be focused on even more streamlined processing and automation to further enhance throughput, particularly of whole mice and 3D-Swiss rolls. Also, more use cases will certainly help the technology to establish ROCKETS a useful tool for preclinical drug development and thereby boost the integration into established work streams.

## 5 Acknowledgements

We thank the members of the Beilhack and Sauer labs for helpful discussions.

## 6 Author Contributions

Conceptualization: JM, AB, TP

Methodology: JM, MD, NO, TP, AB

Investigation: JM, NO, FO, AG, ML, TP, AB

Visualization: JM, AKW

Supervision: JM, AB, TP, SC, FH, PU, MS, MD

Writing—original draft: JM

Writing—review & editing: JM, AB, NO, TP

## 8 Data Availability Statement

All datasets generated and analyzed for this study can be made available upon request by the corresponding authors.

## 9 Abbreviations

AAALAC: association for assessment and accreditation of laboratory animal care
BABB: 1:2 (v/v) mixture of benzyl alcohol and benzyl benzoate
BLI: bioluminescence imaging
CBD: common bile duct
CP: choroid plexus
CT: computed tomography
DCM: dichloromethane
dH2O: deionized water
DMSO: dimethylsulfoxide
DP: duodenal papilla
ECHA: european chemicals agency
EDTA: ethylenediaminetetraacetic acid
EpCAM: epithelial cell adhesion molecule
EtOH: ethanol
GI tract: gastrointestinal tract
H&E: hematoxylin and eosin
LN: lymph node
LSFM: light sheet fluorescence microscopy
MeOH: methanol
MRI: magnetic resonance imaging
MSG: mucous salivary gland
NBF: neutral buffered formalin
PBS_PC_: phosphate buffered saline (subscript PC: added biocide ProClin300)
PET: position electron tomography
PPs: Peyer’s patches
RI: refractive index
ROCKETS: rapid optical clearing kit for enhanced tissue scanning
RT: room temperature
SSG: serous salivary gland
STO: stomach
SI: small intestine
CAE: caecum
COL: colon
T-80: Tween-80®
TAA: tumor-associated antigens
TEA: triethanolamine
UM: ultramicroscope
v/v: volume per volume (%)
w/v: weight per volume (%)
UM: ultramicroscope

