## Supplemental Material for "ROCKETS - a novel one-for-all toolbox for light sheet microscopy in drug discovery"

*Supplementary Material*

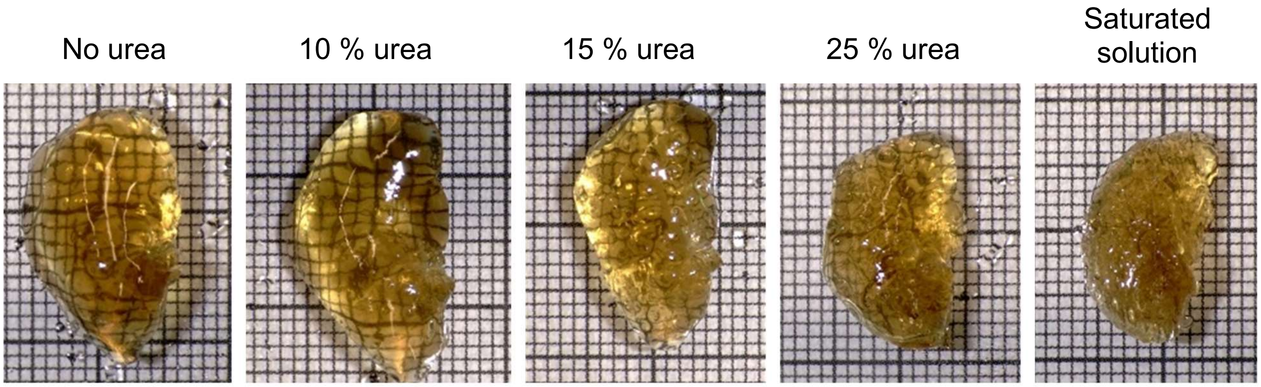


**Supplementary Figure S1. Cleared whole mouse liver lobe specimens after incubation with ROCKETS at indicated concentrations of urea.** Urea concentrations of 15% (w/v) or higher induced deformation of liver (shown: left lateral lobe) and other tissues that is not reversed upon dehydration and clearing. Thick grid lines = 1 cm.

**Supplementary Table S1. Channel usability for whole-organ imaging of non-perfused murine specimens with LSFM after ROCKETS preclearing or untreated.**
Sm./L. int. = Small/Large intestine; SR = Swiss Roll; Fem. repr. = Female reproductive tract. Channel usability is defined as:

✓ = full-depth high-contrast imaging at homogenous trans-illumination

(✓) = full-depth high-contrast imaging, inhomogeneous trans-illumination with brightness gradient towards center

× = imaging of entire sample not possible using the respective channel (blurry or dark center)

| | ROCKETS precleared | | | | | Untreated (PBS) | | | | |
|---|---|---|---|---|---|---|---|---|---|---|
| **Channel** | **1** | **2** | **3** | **4** | **5** | **1** | **2** | **3** | **4** | **5** |
| **Ex. λ [nm]** | **470** | **545** | **630** | **685** | **747** | **470** | **545** | **630** | **685** | **747** |
| **Spectrum** | | | | | | | | | | |
| Kidney | (✓) | ✓ | ✓ | ✓ | ✓ | × | × | (✓) | ✓ | ✓ |
| Tongue | ✓ | ✓ | ✓ | ✓ | ✓ | (✓) | ✓ | ✓ | ✓ | ✓ |
| Heart | (✓) | ✓ | ✓ | ✓ | ✓ | × | × | × | ✓ | ✓ |
| Liver | (✓) | ✓ | ✓ | ✓ | ✓ | × | × | × | × | (✓) |
| Spleen | (✓) | ✓ | ✓ | ✓ | ✓ | × | × | (✓) | ✓ | ✓ |
| Stomach | ✓ | ✓ | ✓ | ✓ | ✓ | (✓) | ✓ | ✓ | ✓ | ✓ |
| Sm. int. (SR) | (✓) | ✓ | ✓ | ✓ | ✓ | × | (✓) | ✓ | ✓ | ✓ |
| L. int. (SR) | ✓ | ✓ | ✓ | ✓ | ✓ | × | (✓) | ✓ | ✓ | ✓ |
| Caecum | ✓ | ✓ | ✓ | ✓ | ✓ | (✓) | ✓ | ✓ | ✓ | ✓ |
| Fem. repr. | ✓ | ✓ | ✓ | ✓ | ✓ | ✓ | ✓ | ✓ | ✓ | ✓ |
| Lungs | ✓ | ✓ | ✓ | ✓ | ✓ | × | (✓) | ✓ | ✓ | ✓ |
| Sal. Glands | ✓ | ✓ | ✓ | ✓ | ✓ | × | × | ✓ | ✓ | ✓ |
| Pancreas | (✓) | ✓ | ✓ | ✓ | ✓ | (✓) | ✓ | ✓ | ✓ | ✓ |
| Lymph node | ✓ | ✓ | ✓ | ✓ | ✓ | ✓ | ✓ | ✓ | ✓ | ✓ |
| Bladder | ✓ | ✓ | ✓ | ✓ | ✓ | ✓ | ✓ | ✓ | ✓ | ✓ |
| Mam. Glands | ✓ | ✓ | ✓ | ✓ | ✓ | ✓ | ✓ | ✓ | ✓ | ✓ |
| Thymus | ✓ | ✓ | ✓ | ✓ | ✓ | × | (✓) | ✓ | ✓ | ✓ |

**Supplementary Table S2. LSFM-based biodistribution scoring of G8.8R binding after intravenous administration (20 µg, 24 h) based on fluorescence intensity levels.**

| Tissue/Structure | Score |
|---|---|
| **Kidneys and urinary tract** | |
| Glomeruli | |
| Bowman's capsule | - |
| Capillary tufts | - |
| Proximal convoluted tubules | - |
| Henle's loops | |
| Descending loop | - |
| Ascending loop | - |
| Collecting ducts* | +++ |
| Medulla† | ++ |
| Pelvis† | + |
| Ureter | + |
| Bladder epithelium | + |
| Urethra | + |
| **Lung** | |
| Tracheal mucosa | + |
| Bronchi | + |
| Bronchioli | ++ |
| Alveolar AT1 cells | - |
| Alveolar AT2 cells | + |
| **Pancreas** | |
| Acinar cells | + |
| Duct cells | + |
| Islet cells | - |
| **Liver** | |
| Hepatocytes | - |
| Gall bladder | |
| Mucosa | + |
| Bile canaliculi | ++ |
| Bile ductules and ducts | + |
| **Skin** | |
| Epidermal Kertinocytes | - |
| Hair bulb and shaft | - |
| Hair root and sheath | - |
| Sebaceous glands | ++ |
| Sweat glands | |
| Acinar cells | ++ |
| Duct cells | +++ |
| Myoepithelial cells | - |
| **Brain‡** | |
| Neurons | - |
| Glia cells | - |
| Brain ependymal cells | - |
| Choroid plexus ependymal cells | ++ |

| Tissue/Structure | Score |
|---|---|
| **Mammary glands** | |
| Ductal epithelium | +++ |
| Alveolar epithelium | - |
| **Salivary glands** | |
| Serous acinar cells | ++ |
| Mucous acinar cells | + |
| Ductal cells | ++ |
| **Oral cavity** | |
| Tongue, squamous epithelium | - |
| Gustatory papillae (foliate, fungiform and circumvallate) | ++ |
| Larynx | ++ |
| **Stomach** | |
| Esophagus tunica mucosa | + |
| Glandular gastric epithelium§ | +/++ |
| Forestomach squamous epithelium | - |
| Limiting ridge epithelium | + |
| Pylorus epithelium§ | +/++ |
| **Small intestine‖** | |
| Villi, epithelial cells | + |
| Crypts, epithelial cells | ++ |
| Duodenal papillae, cmn. bile duct | +++ |
| Peyer's patch, adjacent epithelium | +++ |
| Peyer's patch, follicles¶ | + |
| **Large intestine (caecum, colon, rectum)** | |
| Crypts, epithelial cells# | ++ |
| Caecal Peyer's patch | ++ |
| **Female reproductive organs** | |
| Uterus and vaginal endometrium | + |
| Cervix endometrium | - |
| Oviducts mucosal epithelium | |
| Ampullae | +++ |
| Isthmus of ampullae and uterus | + |
| Infundibulum | + |
| Ovaries** | |
| Germinal epithelium | + |
| Ovarian bursa and follicles | - |
| **Thymus** | |
| Medullary thymic epithelium | ++ |
| Cortical thymic epithelium | + |
| **Connective, adipose and muscular tissues; skeletal bones** | - |
| **Lymphoid organs (lymph nodes, spleen, Peyer's patch follicles)** | (+) |
| Non-identified signals | |

*Cortical, medullary, papillary ducts (ducts of Bellini). Associated intercalated cells showed increased binding. Binding generally higher for cortical nephrons than juxtamedullary nephrons.
†Individual structures in the medulla and pelvis not identified due to lack of contrast.
‡Brains were not precleared and dehydrated/delipidated using MeOH/DCM, which may have affected fluorescence signals differently than preclearing with EtOH-dehydration (as conducted for all other tissues)
§ Glandular stomach highly heterogeneous binding pronounced near limiting ridge, lesser curvature
|| Binding levels and pattern equal along duodenum, jejunum and ileum.
¶ Signal pattern in lymphoid organs was non-polarized and likely originates from immunogenicity of the antibody and corresponding reactions of the hosts immune system.
### General gradient of binding pronounced towards crypt base/decreased towards luminal surface.
**We detected an unidentified spherical structure within the ovarian bursa that was highly positive but could not be allocated to any normal ovarian cell type (compare Suppl. Fig. S14E and F).

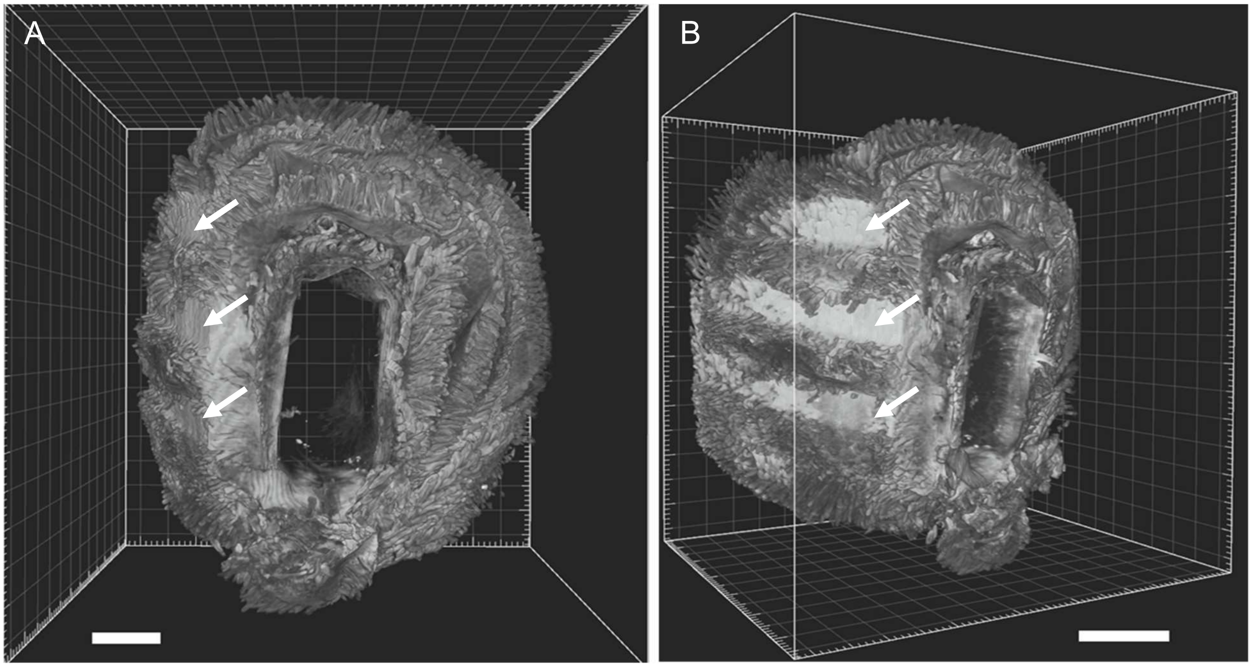


**Supplementary Figure S2. Deformation artifact of a 3D-Swiss Roll of the small intestine through contact with a processing cassette during fixation. (A)** Top view of a specimen of the small intestine (SI 1) that was squeezed and flattened (left side) by being pressed to the surface of a sample-processing cassette (arrows). **(B)** Side view of the same specimen with visible slits of the histology cassette imprinted in the tissue. Scale bars = 1 cm.

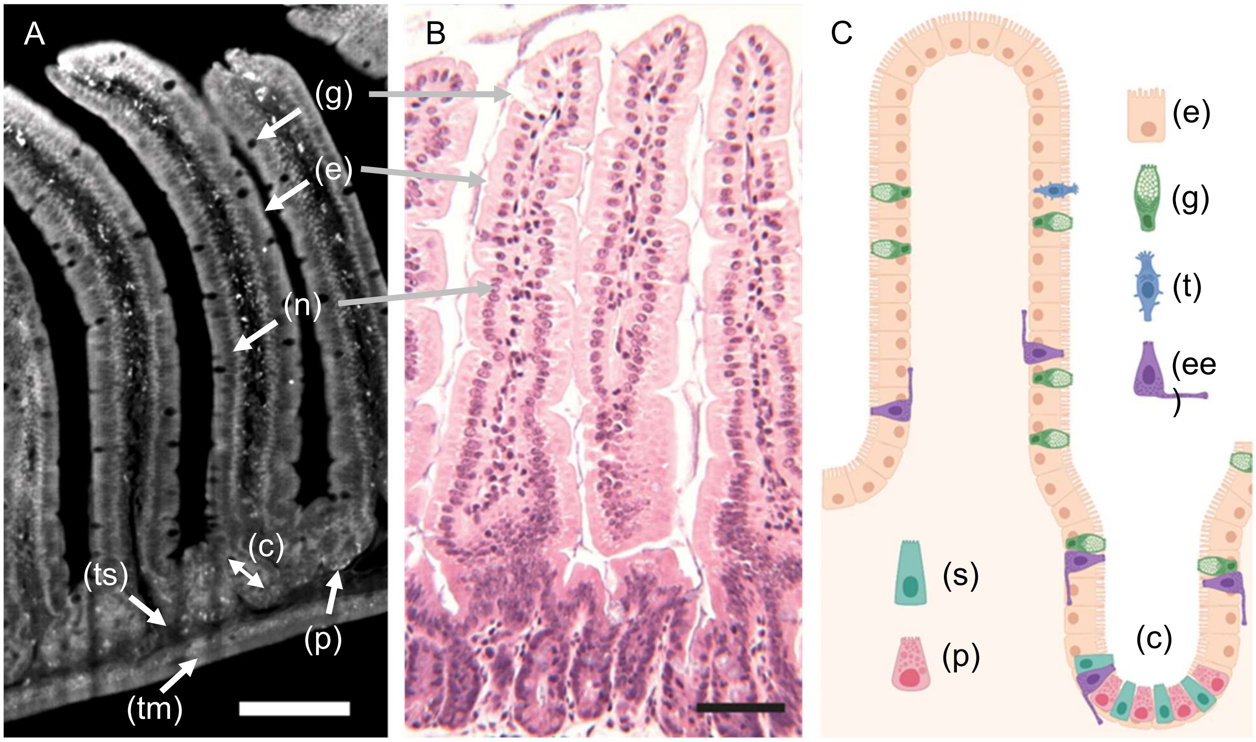


**Supplementary Figure S3. Light Sheet Fluorescence Microscopy (LSFM) enables analysis of microanatomical features and identification of cell types. (A)** Autofluorescence (grey) in a single high-resolution image of an LSFM scan of a 3D-Swiss Roll of the small intestine showing intact microanatomical tissue features. **(B)** Photomicrograph of a physical section of the small intestine stained with hematoxylin and eosin (H&E) as shown for a standard preparation technique (*49*). **(C)** Diagram of the murine small intestine depicting one villus with crypt and intestinal cell types. Enterocyte (e), nuclei (n), goblet cell (g), tuft cell (t), enteroendocrine cell (ee), stem cell (s), paneth cell (p), crypts of Lieberkühn (c), tela submucosa (ts), tunica muscularis (tm). Scale bars = 100 µm.

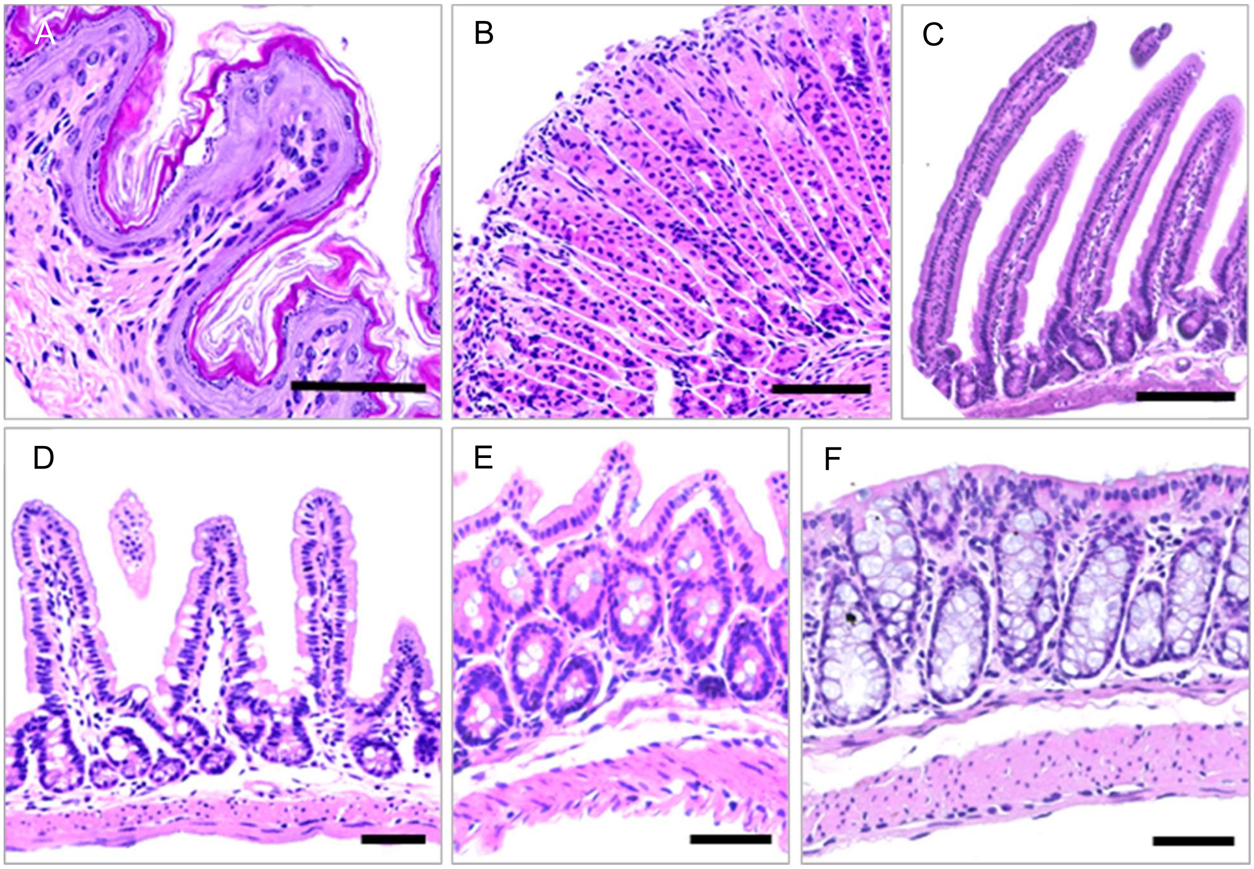


**Supplementary Figure S4. Physical sections of the GIT stained with H&E following 3D-Swiss Roll processing, dehydration, clearing and LSFM imaging. (A)** Forestomach **(B)** Glandular stomach fundus **(C)** Duodenum **(D)** Ileum **(E)** Caecum **(F)** Colon. All specimens show regular microanatomical features and staining characteristics after processing as 3D-Swiss Rolls, clearing and LSFM imaging. Scale bars = 50 µm.

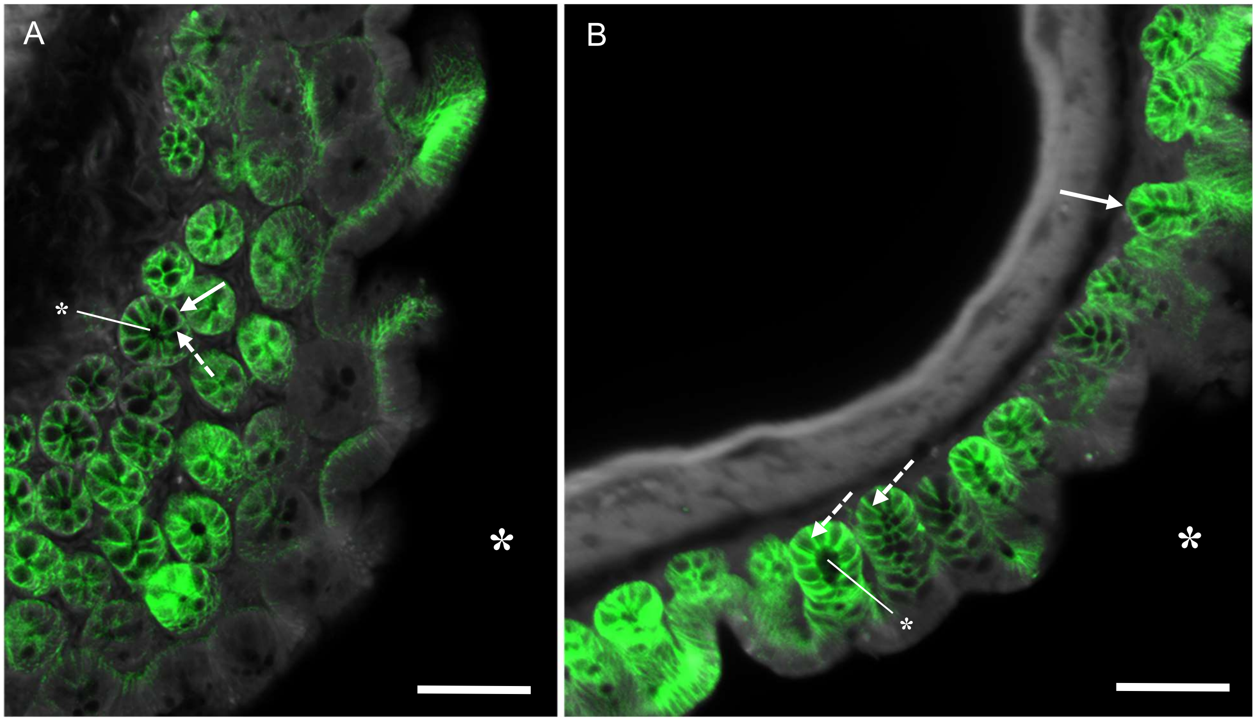


**Supplementary Figure S5. Single LSFM images reveal basolateral EpCAM--AF750-binding patterns to epithelia. (A)** Caecum and **(B)** colon with polarized binding patterns of the antibody to lateral (dashed arrows) and basal (arrows) membranes of epithelial enterocytes. This binding pattern was observed for all normal simple epithelia throughout the body. *Luminal side of the tissue is indicated by an asterisk. Scale bars = 50 µm.

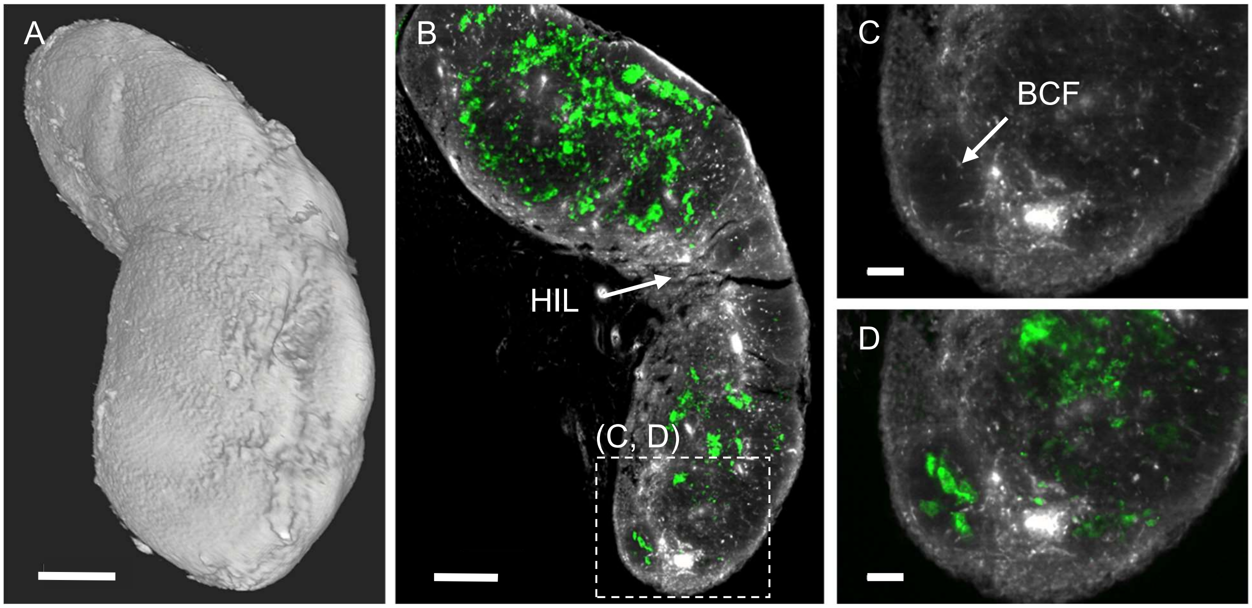


**Supplementary Figure S6. Anti-EpCAM-AF750 antibody staining in the lymph nodes. (A)** 3D-Surface rendering and **(B)** Maximum intensity projection of a z = 100 µm virtual section ($MIP_{100µm}$) of a single inguinal lymph node at approximately half width (central) depicting general anatomy (grey) and antibody binding (green). **(C, D)** Single digital sections of the area indicated in image (B). Hilus (HIL), B cell follicle (BCF, *bona fide*). Scale bars = 200 µm (A, B), 50 µm (C, D).

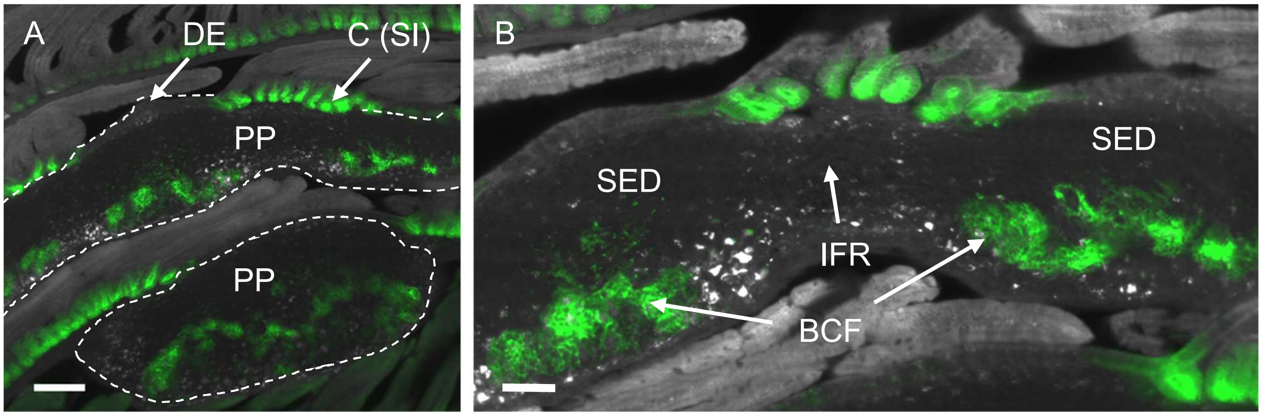


**Supplementary Figure S7. Anti-EpCAM-AF750 antibody binding in Peyer's patches (PP).** **(A)** $MIP_{100\mu m}$ of two PPs (encircled) in consecutive layers of the 3D-Swiss Roll of a small intestine (see also Fig. 6.41, D). Note the increased antibody binding to crypts of the small intestine C(SI) between two follicles of one PP. DE = dome epithelium. **(B)** $MIP_{100\mu m}$ of the upper PP (encircled) depicted in image (A), B cell follicles (BCF*, bona fide*) with antibody binding (green). Note the decreasing signal intensity at the DE. The subepithelial dome (SED) and interfollicular regions (IFR) were always excluded from antibody binding. Scale bars = 200 µm (A), 50 µm (B).

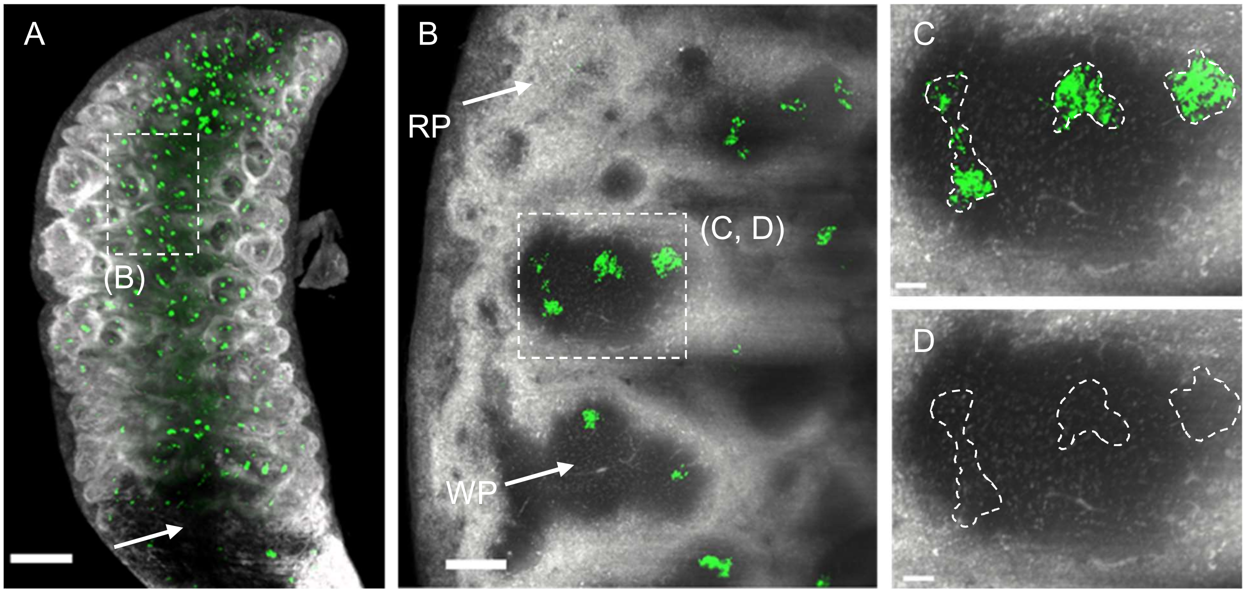


**Supplementary Figure S8. Anti-EpCAM-AF750 antibody binding in the spleen. (A)** $MIP_{WHOLE}$ of the entire spleen anatomy (grey) and antibody binding (green). Note melanosis on the bottom left of the organ (arrow). **(B)** $MIP_{100\mu m}$ of the region indicated in image (A) depicting several white pulp areas (WP, darker spots in autofluorescence, grey), red pulp (RP, bright areas) and the capsule (edge). **(C, D)** $MIP_{100\mu m}$ of the region indicated in image (B). Note that antibody accumulations (green) do not correspond to any distinct structures in the autofluorescence channel (encircled, D). Scale bars = 1000 μm (A), 150 μm (B), 30 μm (C, D).

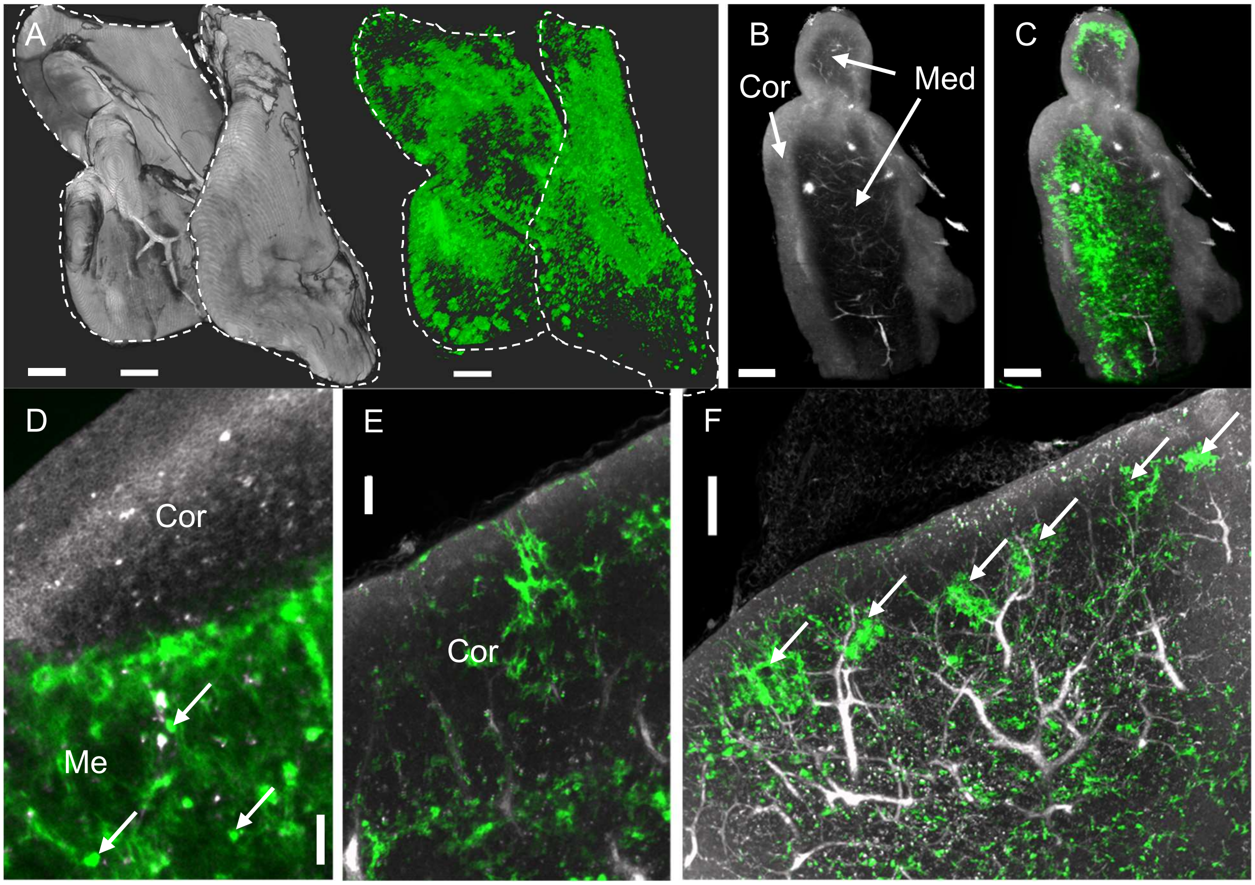


**Supplementary Figure S9. Anti-EpCAM-AF750 antibody binding in in the thymus. (A)** Surface rendering of both lobes (each encircled) of the thymus (left, grey) and respective antibody binding (right, green). **(B)** $MIP_{200\mu m}$ of the left lobe anatomy (grey, Ch2) and **(C)** anatomy and antibody tissue binding as overlay (green, Ch5). **(D)** Single LSFM image depicting the interface of cortex (Cor) and medulla (Med) with locally restricted antibody binding in the medulla. Arrows indicate some of many speckles detected within the overall mesh-like binding pattern. **(E)** Another region of the same specimen showing binding also in the cortex. **(F)** $MIP_{100\mu m}$ of the thymus showing focally increased binding in distinct areas (arrows). Scale bars = 250 µm (A-C), 40 µm (D), 50 µm (E), 100 µm (F).

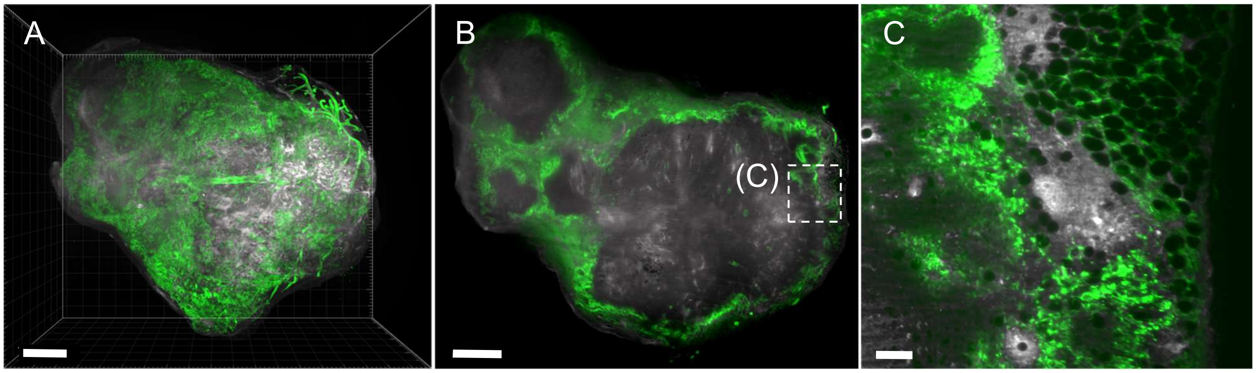


**Supplementary Figure S10. Anti-EpCAM-AF750 antibody binding in the tumor. (A)** 3D-rendering of a subcutaneous tumor of a pancreatic cancer cell line (KPC-4662, green) **(B)** Single digital section (slice 527/661) of the tumor depicting highly heterogeneous binding of the antibody. **(C)** Higher magnification (10x) image of the region indicated in (B). Scale bars = 1000 µm (A, B), 100 µm (C)

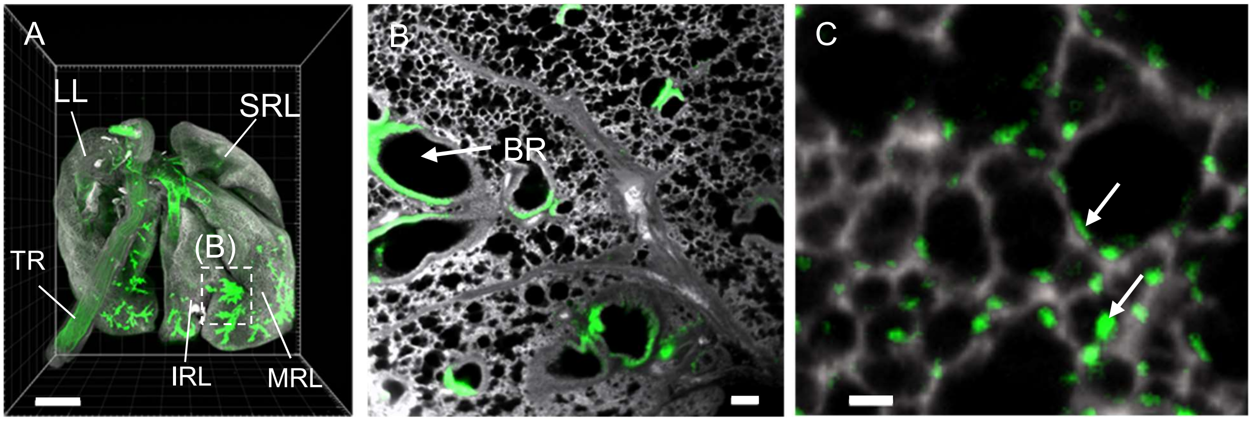


**Supplementary Figure S11. Anti-EpCAM-AF750 antibody binding in in the lower respiratory tract. (A)** Dorsal view of the gross lung anatomy (grey) and antibody binding (green). Left lobe (LL), superior right lobe (SRL), medial right lobe (MRL), Inferior right lobe (IRL), trachea (TR). View of post-caval lobe is obstructed. **(B)** Enlarged single slice view of the region indicated in (A) depicting several bronchioli (BR) and detected antibody (green). **(C)** Scattered localization of the antibody within the alveoli protruding into the alveolar space (arrows). Scale bars: 1000 µm (A), 200 µm (B) and 100 µm (C).

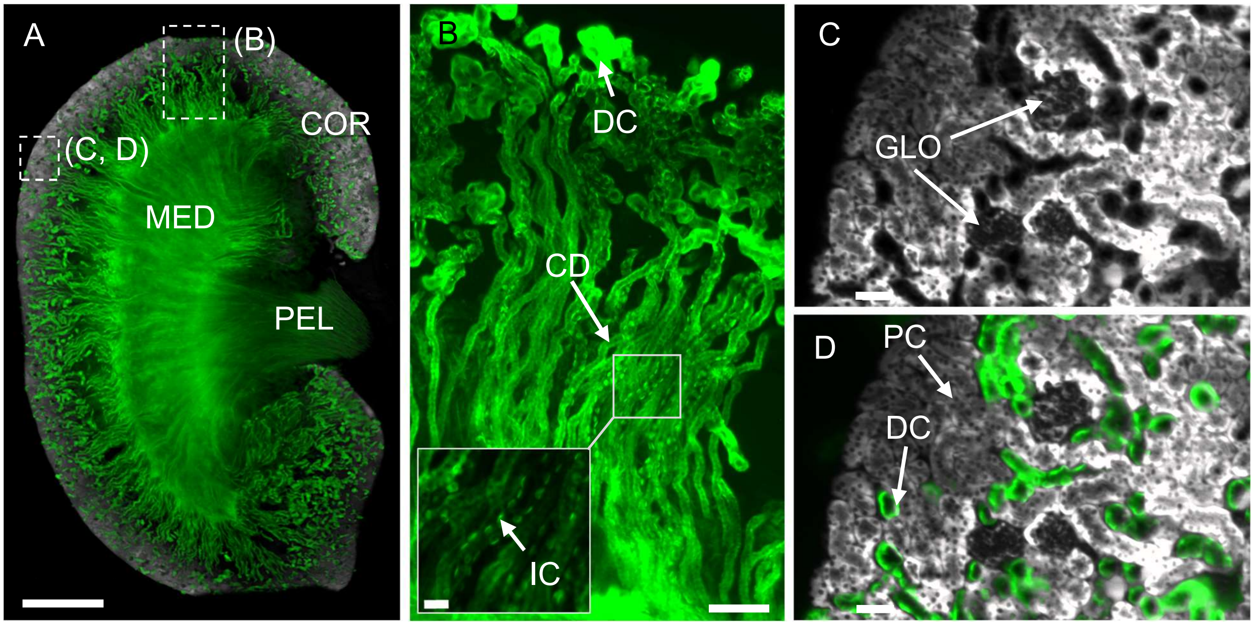


**Supplementary Figure S12. Anti-EpCAM-AF750 antibody binding in the kidney. (A)** $MIP_{50\mu m}$ at the center of the kidney depicting its anatomy (grey) and bound antibody (green). Medulla (MED), Cortex (COR), Pelvis (PEL). **(B)** $MIP_{30\mu m}$ of area indicated in (A) depicting only the antibody signal. Distal convoluted tubule (DC), proximal convoluted tubule (PC), collecting duct (CD), intercalated cell (IC). Scale bars: 2000 µm (A), 10 µm (B, box), 50 µm (B, C and D).

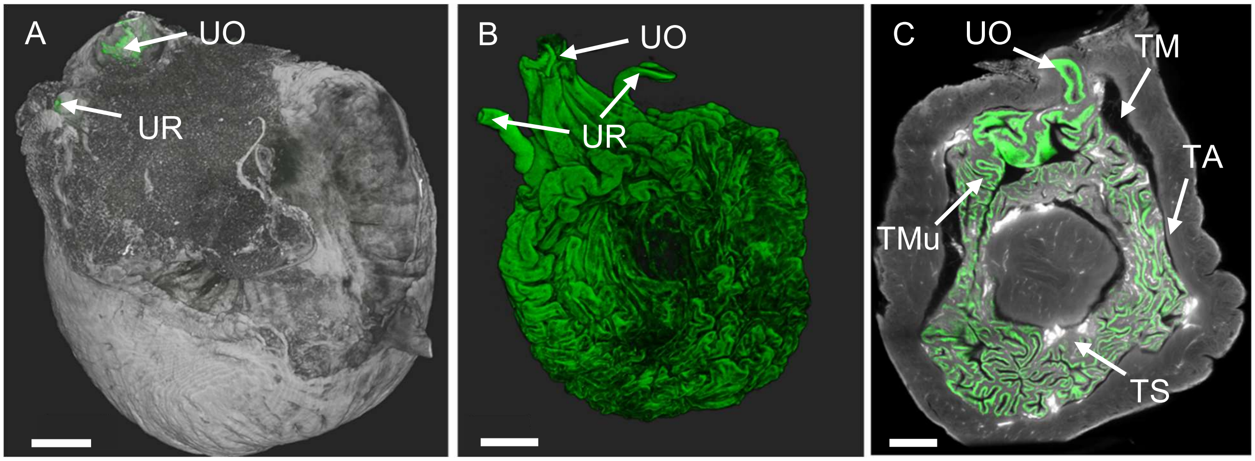


**Supplementary Figure S13. Anti-EpCAM-AF750 antibody binding in the bladder (A)** Rendering of the entire bladder anatomy **(B)** Surface rendering of the antibody binding in the entire bladder. **(C)** Single LSFM image depicting the bladder anatomy (grey) and bound antibody (green). Urethra (U), urethral opening (UO), tunica adventitia (TA), tunica subserosa (TS), tunica muscularis (TM), Tunica mucosa (TMu). Scale bars = 400 µm.

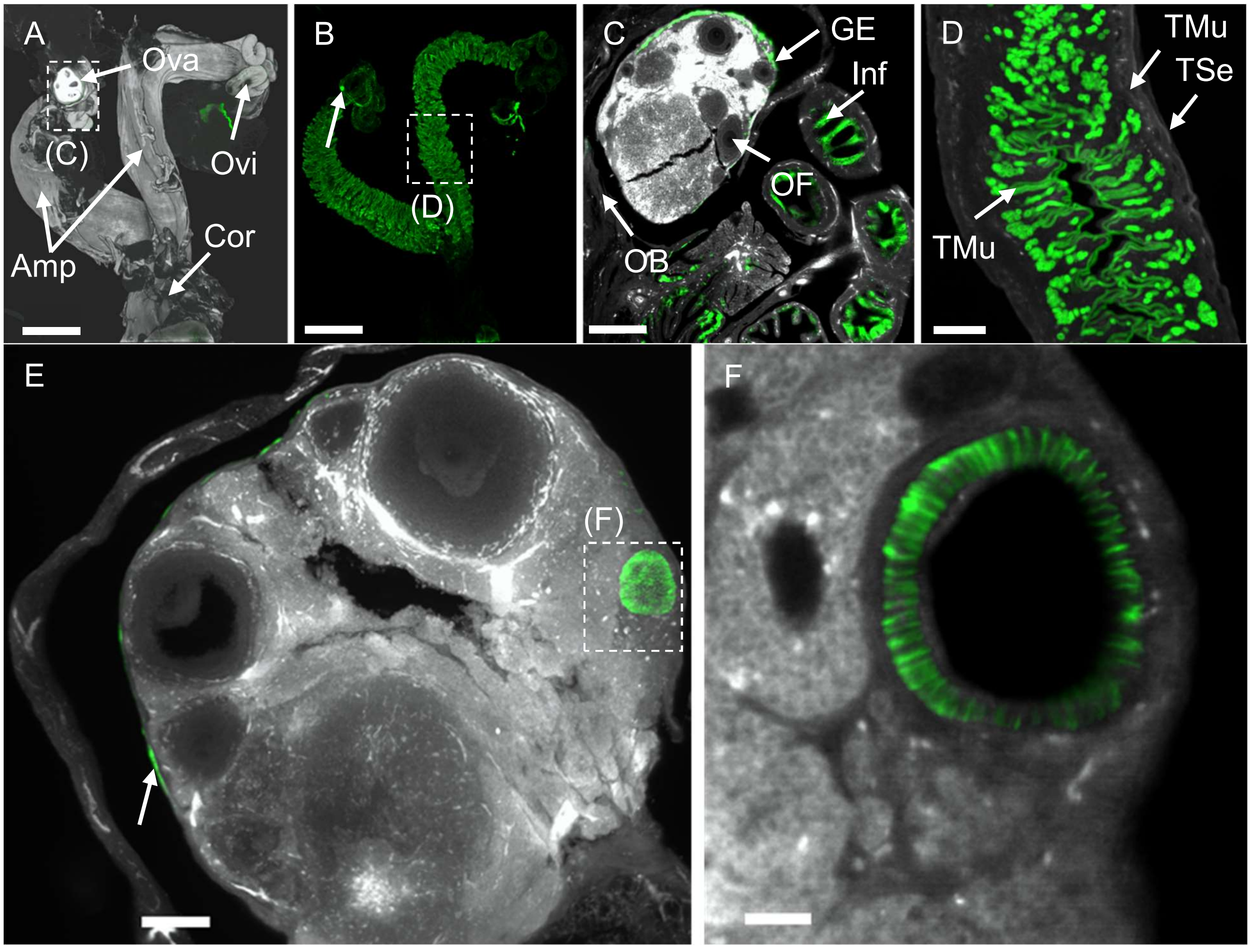


**Supplementary Figure S14. Anti-EpCAM-AF750 antibody binding in the female reproductive organs**. **(A)** General morphology of the specimen (grey) and **(B)** overview of EpCAM tissue binding (green). Arrow indicates a spherical accumulation of antibody in one of the ovaries (see also E, F). **(C)** Single LSFM image of one of the ovaries and oviducts as indicated in (A). **(D)** Single LSFM image of one of the ampullae as indicated in image (B). Corpus (Cor), infundibulum (Inf), ovary (Ova), oviduct (Ovi), tunica mucosa (TMuc, endometrium), tunica muscularis (TMus), tunica serosa (TSer), ovarian follicle (OF), ovarian bursa (OB), germinal epithelium (GE), ampulla (Amp). **(E)** $MIP_{500\mu m}$ of the left ovary showing antibody binding to the germinal epithelium (arrow) and an unidentified spherical object with very high antibody binding (boxed, and F). **(F)** Single Image as indicated in (E) showing lateral membranous binding. Scale bars = 1000 µm (A, B), 300 µm (C), 200 µm (D), 100 µm (E), 30 µm (F).

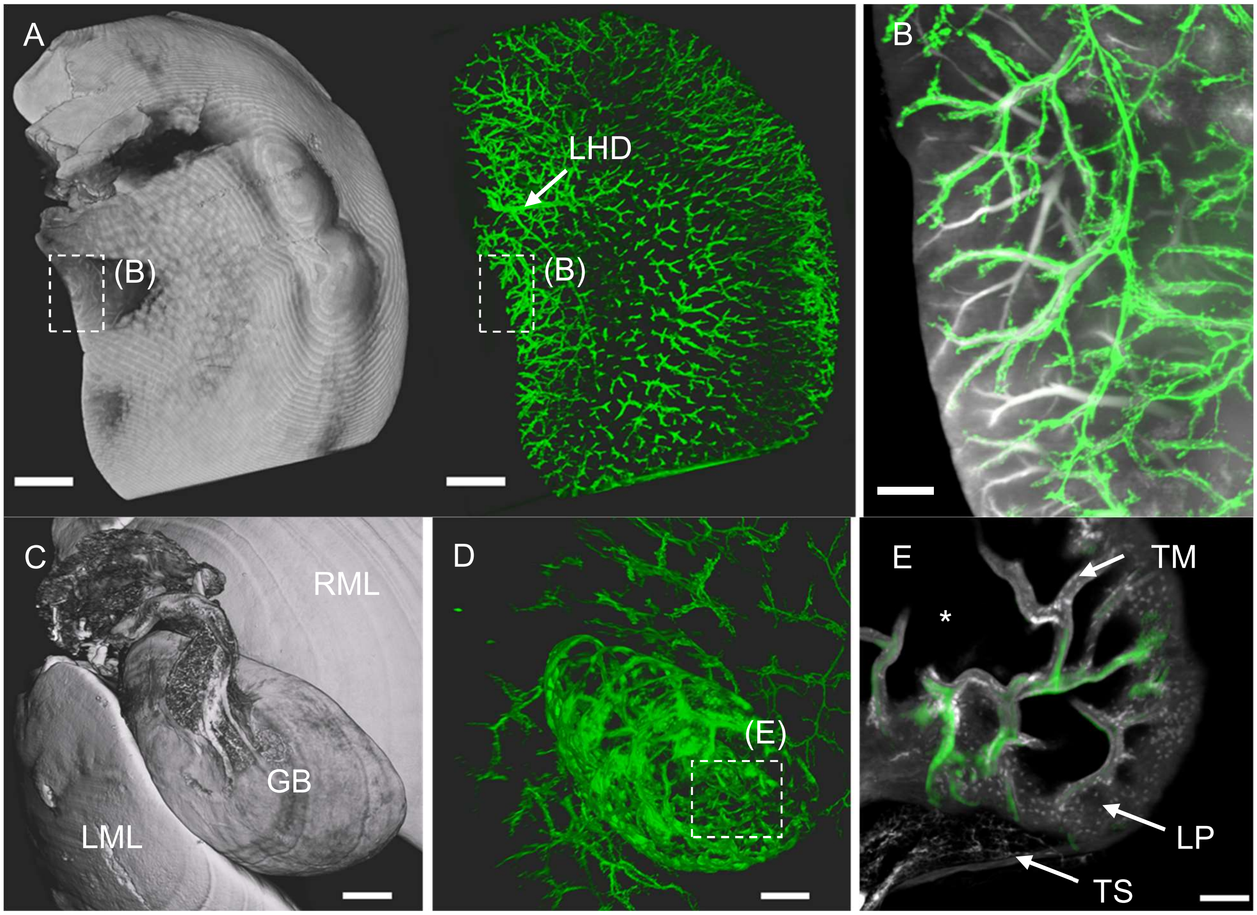


**Supplementary Figure S15. Anti-EpCAM-AF750 antibody binding in the liver and gall bladder. (A)** Ventral surface rendering of left liver lobe anatomy (left, grey) and $MIP_{WHOLE}$ of antibody (right, green) bound to the entire biliary ductal tree with visible left hepatic duct (LHD). **(B)** $MIP_{200\mu m}$ overlay of both channels as indicated in (A). Antibody binding to biliary ducts was closely associated with blood vessels. **(C)** Surface rendering of the gall bladder (GB) anatomy *in situ* situated between left (LML) and right (RML) medial liver lobe. **(D)** Antibody binding as detected in (C) bound to the GB and biliary ducts in the adjacent liver lobes. **(E)** $MIP_{50\mu m}$ of the gall bladder fundus as indicated in (D) with visible lamina propria (LP), tunica serosa (TS) and tunica mucosa (TM). *asterisks indicates luminal side. Scale bars = 2000 µm (A), 200 µm (B), 300 µm (C, D), 50 µm (E).

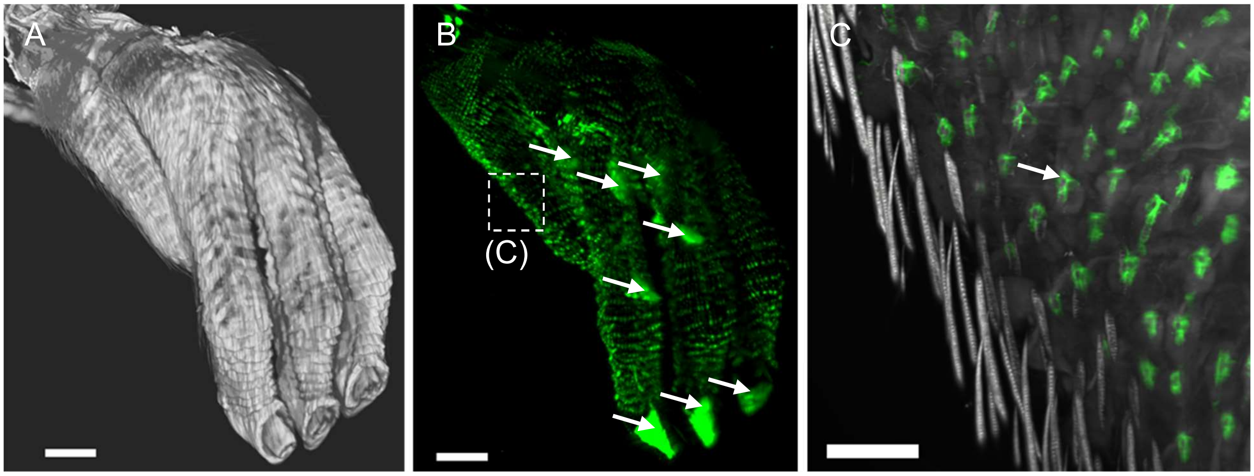


**Supplementary Figure S16. Anti-EpCAM-AF750 antibody binding at hair follicles and sweat glands in the paw.** **(A)** 3D-surface rendering of the anatomy of the left paw. **(B)** $MIP_{WHOLE}$ of the antibody binding with increased binding to all footpads (arrows). **(C)** $MIP_{50\mu m}$ of the general tissue anatomy (grey) and bound antibody as indicated in (B). Arrow indicates one single hair follicle with antibody bound at sebaceous glands. Scale bars = 1000 µm (A, B), 200 µm (C).

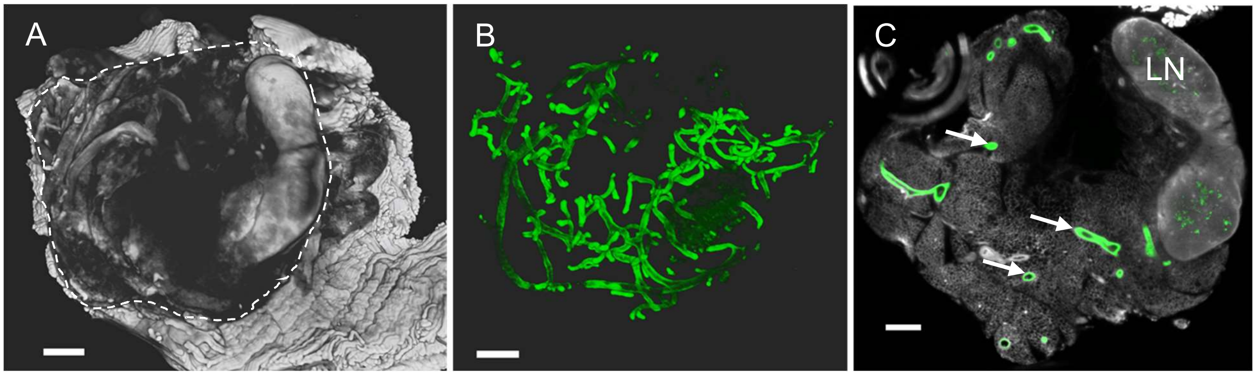


**Supplementary Figure S17. Anti-EpCAM-AF750 antibody binding in the mammary glands.** **(A)** 3D-surface rendering of a mammary gland embedded in muscular and adipose tissue. **(B)** 3D-surface rendering of the bound antibody at the lactiferous ducts. **(C)** Single slice view depicting antibody binding (green) within the mammary gland tissue at the epithelium of the ducts (arrows). Note the lymph node (LN) that was by-sampled together with the glandular tissue. Scale bars = 200 μm.

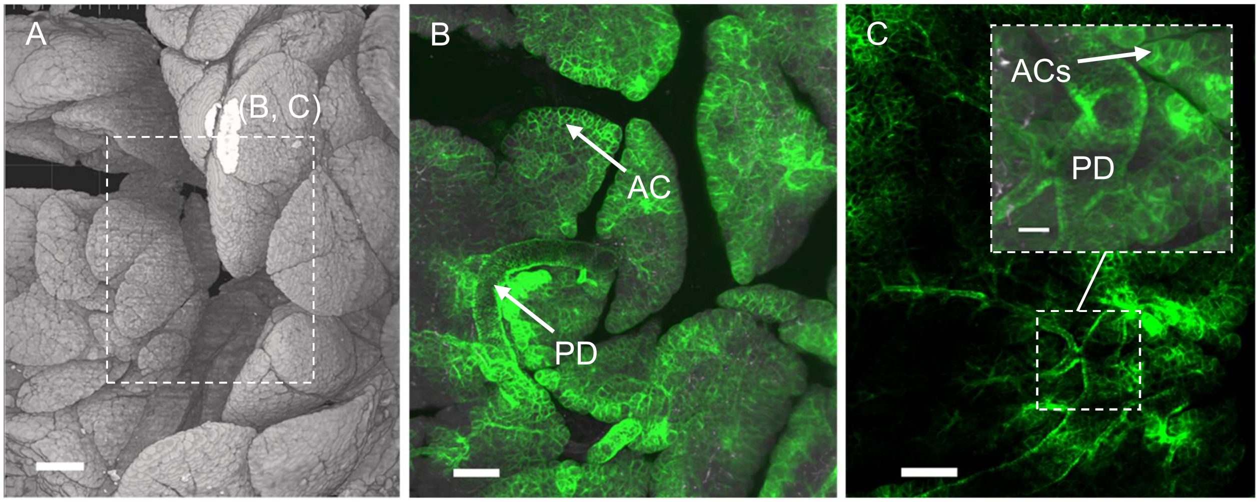


**Supplementary Figure S18. Anti-EpCAM-AF750 antibody binding in the pancreas (A)** Partial 3D-surface rendering of the gastric division of the pancreas. **(B)** MIP$_{100\mu m}$ of the area indicated in (A) showing the general tissue anatomy (grey) and antibody binding (green). Binding is restricted to membranes of acini (AC) and epithelia of pancreatic ducts (PD). **(C)** MIP$_{100\mu m}$ of the same area as image (B) at a different depth with Ch2 (autofluorescence) turned off. Enlargement depicts a branched pancreatic duct (*PD*) with antibody bound evenly distributed to the entire epithelium and surrounding positive acinar cells (*ACs*). Scale bars = 200 µm (A and B), 50 µm (C).
